# Control of Guanine Exchange Factor Activity Using *De Novo* Designed Protein Inhibitors

**DOI:** 10.64898/2026.05.17.725778

**Authors:** Fabiola Vacca, Daniel J. Marston, Caitlin Harris, Pranav Kannan, Heman Burre, Jayani Christopher, Irina Dumbravanu, Mihai L. Azoitei

## Abstract

Guanine Exchange Factors (GEF) of the Dbl family are the main activators of RhoA GTPases. GEF and GTPase activity is tightly regulated at the subcellular level with fast kinetics. Therefore, to fully understand the function of Dbl GEFs requires their study in living cells. Towards developing molecular tools that reversibly and rapidly modulate the activity of endogenous GEFs in living cells, here we developed a general platform for engineering inhibitors against members of the Dbl family of GEFs using generative protein design. Engineered proteins showed high affinity and remarkable specificity for the target GEFs and modulated GEF activity both *in vitro* and in cells. In a proof-of-principle example, a GEF inhibitor was coupled to a light-activated module, enabling the optogenetic control of its activity in cells. These findings show that generative protein design can create modulators of intracellular signaling and broaden the range of tools available for biological research.

## Introduction

Cellular processes are governed by complex signaling networks that are precisely coordinated in space and time. Such spatiotemporal control is essential for cell migration^1^, proliferation^2^, and immune recognition^3^. Rho GTPases and their activators, guanine nucleotide exchange factors (GEFs), play a central role in these processes by regulating cytoskeletal changes. Cdc42, Rac1, and RhoA are well characterized Rho GTPases that are master regulators with distinct roles in actin cytoskeleton remodeling: Cdc42 promotes protrusive filopodia, RhoA induces contractile actin-myosin filaments, and Rac1 stimulates the formation of protrusive lamellipodia^4,5^. These morphological changes depend on the timely activation of GTPases by GEFs, which catalyze the exchange of GDP for GTP^6^.

The Dbl family of GEFs has 69 members and constitutes the largest class of Rho GTPase activators. All Dbl GEFs contain a DH domain, which is the site of GTPase interaction, and a PH domain with a less conserved role, including membrane localization or providing additional GTPase contacts. Dbl GEFs are typically autoinhibited, as illustrated by Vav1, a canonical member of this family with well understood structural and functional properties. In Vav1, a helical autoinhibitory domain (AID) covers the GTPase interaction site on the DH domain and requires phosphorylation by Src kinase to become active. While autoinhibition is common, Dbl GEF activity is regulated by diverse activators and inhibitory mechanisms. For example, GEF-H1, another Dbl GEF, is inactive when bound to microtubules and only becomes active upon release and Src phosphorylation. In the GEF Intersectin-2, the DH domain is autoinhibited by intramolecular binding of the SH3 region to the C-terminal portion of the DH domain, blocking Cdc42 binding^7,8,9^.

Recently, FRET-based biosensors were developed by us and others for some Dbl GEFs by inserting fluorophore pairs into the hinge region connecting the DH domain to the AID to capture activation/inhibition dynamics across different functional states. Live cell microscopy of these biosensors revealed, in unprecedented detail, that GEF and GTPase activities are precisely coordinated at the subcellular level with second-level kinetics to modulate the cytoskeleton and cell morphodynamics, highlighting that GEF/GTPase signaling can be best understood by measurements in living cells. While these fluorescent biosensors can provide a wealth of information on the timing and distribution of GEF activity, there are limitations related to extending their design principles to monitor the activity of other GEFs as well as to the type of data they can reveal. The development of GEF biosensors involved extensive empirical testing and required detailed knowledge of the inhibition and activation mechanisms of the target GEF, knowledge which is missing for most Dbl GEFs. Biosensors provide only correlational data between molecular activity and cellular activity, and their overexpression can cause biological artifacts. To develop next generation tools for detecting and manipulating GEF activity, here we describe a general platform to engineer novel proteins that bind with high affinity and specificity to the DH domain of target GEFs and prevent GTPases interaction, acting as inhibitors. These proteins can be functionalized with optogenetic modules or fluorescent probes to control and report on the activity of endogenous GEFs in living cells.

The development of GEF inhibitors has proven challenging because their GTPase binding site is a flat surface that lacks unique features and binding pockets. Additionally, the DH domain region that interacts with GTPases is highly conserved across the 69 Dbl GEFs, making it challenging to target a specific GEF without also affecting the activity of related family members. To date, only one small molecule inhibitor has been reported for the GEF LARG, but its affinity is low, and it cross-reacts with multiple other GEFs, lacking specificity^10,11^. To address these limitations, we employed recently developed Machine Learning protein design methods^12–16^ to engineer protein binders against the DH domains of Dbl GEFs, Vav1, and Intersectin-2. These novel proteins bound the target GEF with high affinity and inhibited their GTPase interactions *in vitro*. Importantly, despite the high sequence homology of the DH domains among the Dbl GEFs, the engineered inhibitors were highly specific to the target GEF. While engineering binders with novel structures have become more accessible, deploying these molecules *in vivo* for biological discovery remains an active area of investigation. To this end, we demonstrated that Vav1 and Intersectin-2 inhibitors regulate GEF activity in living cells, opening the way to apply these tools *in vivo*. For finer control of the localization and timing of GEF activity, one inhibitor was coupled to a light activated module, which allowed its precise release in living cells. Overall, this study provides novel tools for controlling the activity of specific Dbl GEFs, built on a protein engineering platform that should be generally applicable to other members of this family. Furthermore, our work illustrates how generative protein design and optogenetic tools can be combined to create modulators of intracellular signaling, resulting in novel tools for biological research.

## Results

### Computational design of Vav1 inhibitors

Vav1 was the first GEF selected for binder design because its activity is precisely controlled at the subcellular level, which supports studies of local inhibition to understand its function, and because previous studies have elucidated the structural mechanism by which Vav1 activity is regulated^17^. In its native conformation, Vav1 is autoinhibited, with a 3-turn alpha helix AID covering the GTPases binding site. The active state is achieved upon phosphorylation by Src kinase of a central Tyr residues located in the AID, as well as of two other tyrosines located upstream that further stabilize the AID interactions. As a synthetic peptide, the endogenous AID has weak affinity for the DH domain^17^, which is insufficient for effective inhibition. To design high affinity Vav1 inhibitors, we tested different computational strategies, including modifying existing proteins or designing ones with novel structure to either include the endogenous AID or to build contacts with the DH domain fully *de novo*. A key consideration was the engineering of additional contacts beyond those provided by AID, resulting in binders with high affinity and specificity.

In the first approach, the Vav1 AID (positions 173-178) was grafted onto existing protein “scaffolds” selected from the Protein Databank using a previously described motif transplantation algorithm in Rosetta^18^(**Fig. 1**). A large library of small monomeric proteins was queried computationally to identify proteins with exposed backbones that matched (backbone RMSD<0.5 Å) the conformation of the AID. These proteins were then mutated to display key AID residues I173, Y174, L177 and M178 and modeled on the Vav DH domain in the orientation dictated by the native AID to ensure productive interactions. This resulted in candidate inhibitors that, on average, had 18% of their residues mutated relative to the parent protein scaffold (Supplementary Table 1). Seven proteins engineered with this approach were selected for recombinant expression and characterization, but they showed limited expression. The sequence of three designs that showed some expression was further optimized outside the grafted AID residues using ProteinMPNN^13^ (**Supplementary Fig. 1A**), a machine learning algorithm that takes a backbone as an input and generates a sequence predicted to fold into the target structure.

**Figure 1.**
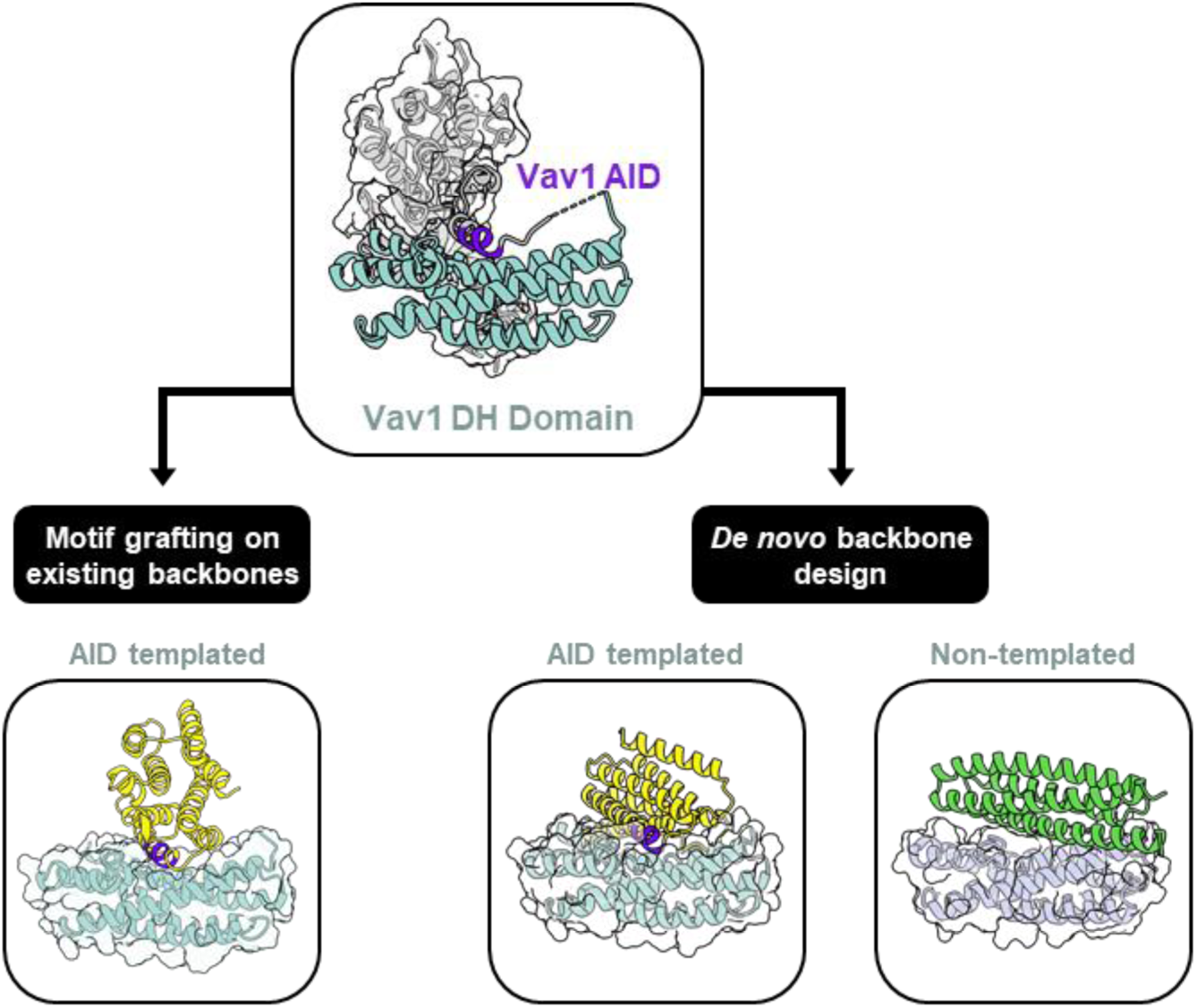
Structure-guided design of Dbl GEF inhibitors. **a.** Schematic overview of the strategies used to design Dbl GEF inhibitors, exemplified by Vav1. The Vav1 CH-DH domain is shown at the center; CH domain: *grey*; Autoinhibitory domain (AID): *purple*; DH domain: *cyan*. *Left*: grafting of the AID backbone into an existing protein (*yellow*). *Right*: *de novo* backbone design, either unconstrained (*green*) or around the AID (*yellow*).

In a second design approach that did not rely on modifying existing proteins, we generated *de novo* binders using a design pipeline that integrated three machine learning algorithms, RFDiffusion^12^, ProteinMPNN, and Alphafold2^14^. In one approach, RFDiffusion designed novel interactions with the DH domain, with contacts guided towards “hotspots” at Vav1 residues 212, 320, 327, and 331 (**Fig. 1**). ProteinMPNN was used to design sequences for the resulting backbones, and AlphaFold-Multimer was used to validate the conformation of the resulting proteins. Out of 11 fully *de novo* designs, 6 were successfully expressed and purified. In another approach, RFDiffusion was guided to design proteins with 40-100 amino acids centered on residues 169-178 of the AID, targeting the same Vav1 “hotspots” as above and focusing on contacting less conserved regions of the DH domain for specificity (**Fig. 1**, also see below). 31 candidate AID-templated inhibitors were selected for experimental characterization, out of which 27 were successfully expressed and purified (**Supplementary Table 2**).

### *In vitro* characterization of Vav1 inhibitors

The inhibitory activity of engineered proteins was evaluated using a fluorescence-based nucleotide exchange assay measuring Vav1 mediated activation of Rac1 via BODIPY-GDP incorporation (**Supplementary Fig. 1B**). The assay consists of pre-incubating the purified DH-PH-CH subunit of Vav1 with the inhibitor, then adding the complex to a reaction containing the GTPase and BODIPY-GDP. Designs that incorporated the Vav1 AID into existing proteins by motif grafting failed to inhibit Vav1 (**Supplementary Fig. 1C**). Similarly, no inhibition was detected for proteins designed fully *de novo* that did not incorporate the Vav1 AID. However, three *de novo* designs built around the AID showed strong Vav1 inhibition, while two others showed weaker activity and were not pursued further (**Fig. 2A,B and Supplementary Fig. 2 D,E**). The three best inhibitors, called Vav1_i6, Vav1_i50, and Vav1_99, were well behaved and stable in solution. Size exclusion analysis revealed that the major species were monomeric and of expected molecular size (**Fig. 2C**), with minimal aggregates or degradation products, a result further confirmed by dynamic light scattering DLS (**Supplementary Fig. 2A**). Differential scanning fluorimetry (DSF) was next used to evaluate the thermal stability of the designs. The Vav1 inhibitors displayed high thermal melting temperatures (Tₘ>70°C), indicative of well-folded, stable proteins (**Supplementary Fig. 2B**).

**Figure 2.**
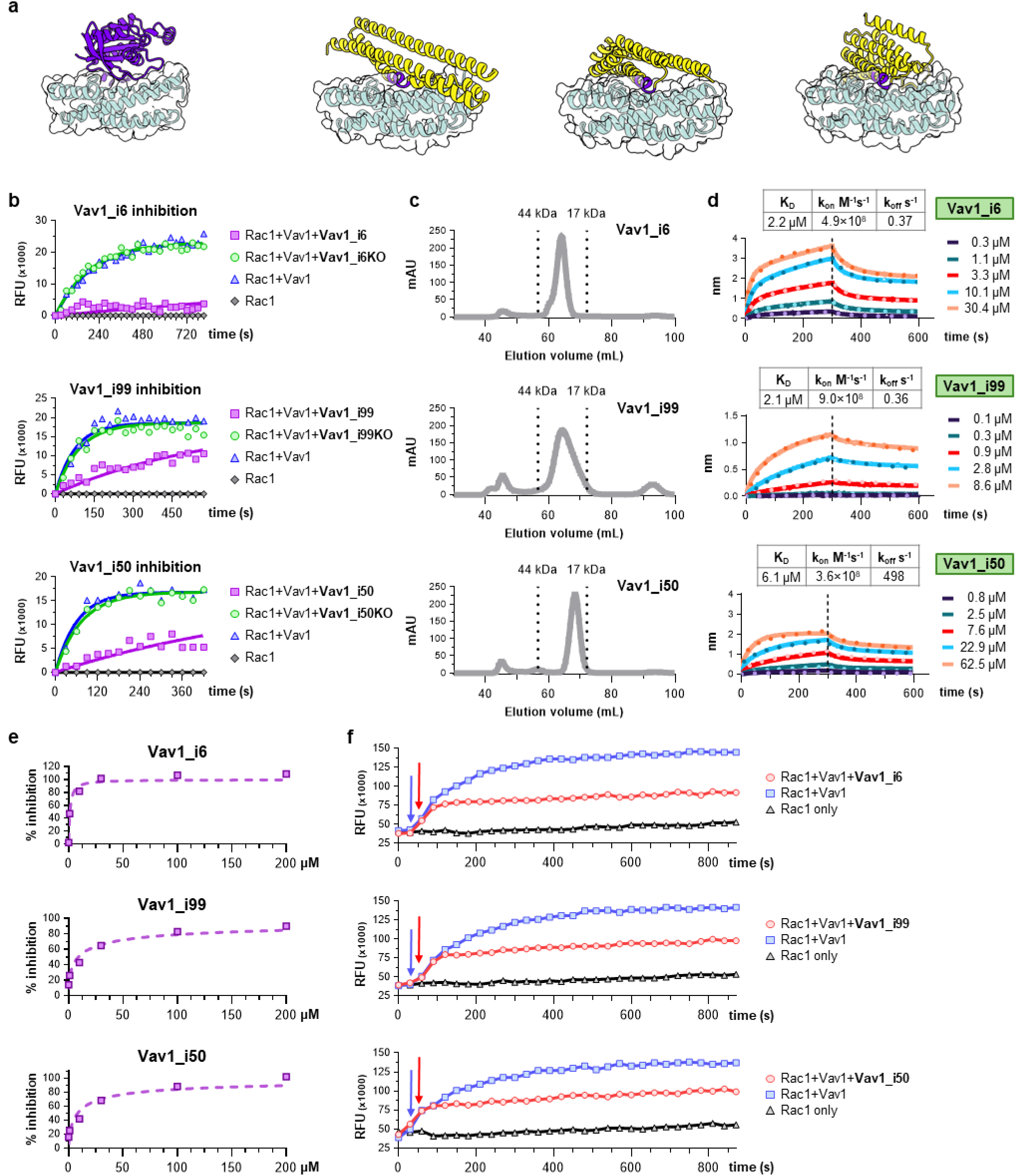
*In vitro* characterization of Vav1 inhibitors. **a.** Structures of Rac1 (*purple*) and engineered inhibitor models in complex with the Vav1 DH domain (*cyan*). **b.** Inhibition off Vav1 exchange activity towards Rac1 by the engineered inhibitors. **c.** Size-exclusion chromatography (SEC) profiles of engineered inhibitors. **d.** BLI binding affinity measurements and of engineered inhibitors to the recombinant VAV1 DH-PH domain. The calculated kinetic parameters (*K*_D_, k_on_, and k_off_) are summarized in the tables after data fitting with a 1:1 binding model. **e.** Concentration–dependent inhibition of Vav1. **f.** Real-time inhibition of showing the inhibition of Vav1 activity upon addition of the inhibitor. Arros indicate addition time for pre-incubated Rac1+ Vav1 +inhibitor (*red*) and Rac1 + Vav1 (*blue*).

Vav1 inhibitors had high affinity for the GEF active domain (DH-PH-CH), with measured equilibrium dissociation constants (*K*_D_s) of 2.1 µM for Vav1_i99, 2.2 µM for Vav1_i6, and 6.1 µM for Vav1_i50, respectively (**Fig. 2D**). The binding was due to predicted contacts, as knock out mutations disrupted Vav1 interactions and limited GEF inhibition (**Supplementary Fig. 2C,D**). To better quantify the inhibition potency, different inhibitor concentrations were tested in the GEF activity assay (**Fig. 2E and Supplementary Fig.,3,4,5**). All inhibitors showed dose-dependent inhibition of Vav1 activity with IC_50_ values of 0.6 μM, 30 μM and 42.2 μM for Vav1_i6, Vav1_i99, and Vav1_i50, respectively. At the maximum concentration tested (60 µM; 300 times higher than the GEF concentration), Vav1_i99 showed a 30-fold reduction in the initial reaction rate (k_obs_) with a 76% decrease in the maximum measured activity over 15 minutes, while Vav1_i6 and Vav1_i50, tested at 40 and 50 µM reduced k_obs_ by 88- and 31 times and the overall GEF activity by 95% and 94% respectively. Full inhibition was also observed in real time by adding the inhibitor to an ongoing GEF/GTPase reaction without GEF/inhibitor pre-incubation (**Fig. 2F**).

### Binding specificity of Vav1 inhibitors

Because the Vav1 inhibitors engage DH domain features that are shared by all 69 members of the Dbl family of GEFs, binding specificity is a potential concern. Structural analysis of the Alphafold2 predictions of the inhibitor designs revealed that while they overlap and occlude the GTPase binding interface of Vav1 through the AID contacts, and thereby disrupt the Vav1 mediated GTPase activation, they also engage discrete DH domain regions that may provide specificity (**Fig. 3A**). Sequence analysis performed on a subset of 12 DH domains of representative Dbl family GEFs shows the conservation of amino acids engaged by the Vav1 inhibitors (**Fig. 3B**). In addition to the core GTPase binding site, which is highly conserved, the inhibitors have interaction sites at the designed hotspots 319/320, which are less well conserved. Furthermore, Vav1_i99 and Vav1_i50 also have additional interaction sites compared to Rac1 at the C-terminus portion of the DH domain (positions 380/1 and 385/6), while Vav1_i6 and Vav1_i99 have interactions at the N-terminus (positions 193 and 197), all of which show high diversity.

**Figure 3.**
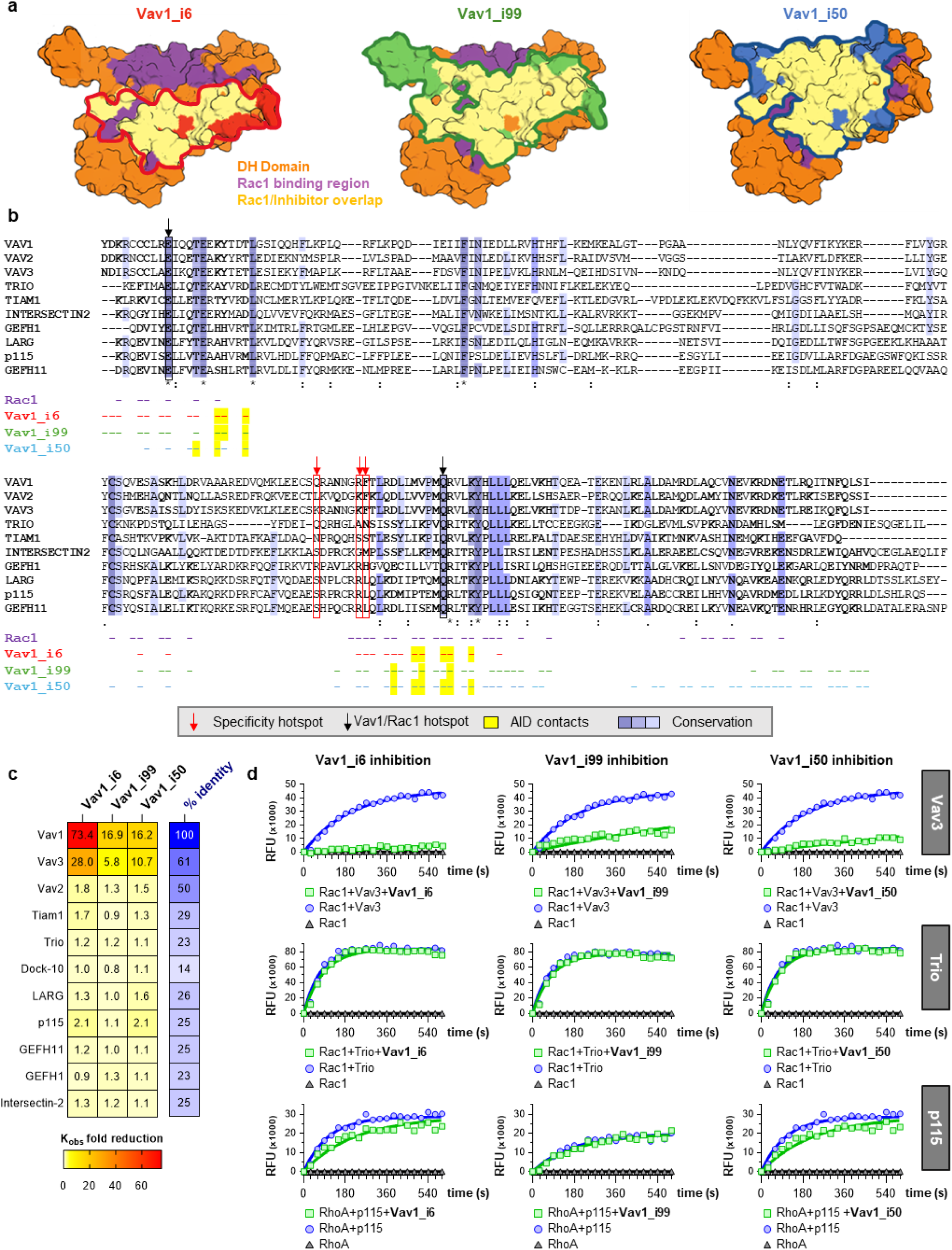
Binding specificity of engineered Vav1 inhibitors. a. Structural mapping of the binding footprint of Rac1 (*purple*) and respective Vav1 inhibitors on the surface of the Vav1 DH domain (*orange*). Binding regions in common between Rac1 and the inhibitors are highlighted in *yellow*. Inhibitor specific contact regions are indicated in *red* (Vav1_inh_6), *green* (Vav1_inh_99), and *cyan* (Vav1_inh_50). b. Sequence alignment of representative Dbl GEFs, with conservation score shown in *purple*. *Red arrows* indicate GEF site targeted by the inhibitor to provide specificity. *Black arrows* indicate Vav1 residues targeted by Rac1 that are also engaged by the inhibitors. Residues in *yellow* highlight DH domain sites engaged by the the Vav1 autoinhibitory domain (AID). c. Fold reduction in the activity of a panel of GEFs by the Vav1 inhibitors. K_obs_ values were calculated from GEF inhibiton assays. The percent sequence similarity between the DH domains of the respective GEFs and Vav1 is shown in *blue*. d. Inhibition data of the Vav1 inhibitors against Vav3, Trio, and p115 Dbl GEFs.

To assess the specificity experimentally, the Vav1 inhibitors were tested against a panel of Dbl family GEFs that target different GTPases: Vav2 and Vav3 (multi-specific); TIAM1, TRIO N-term, DOCK10, and FARP1 (Rac1 GEFs); ARHGEF11, LARG, p115, GEFH1 (RhoA GEFs); and Intersectin-2 (Cdc42 GEF). These GEFs have sequence homology ranging from 25 to 61% with Vav1 across the catalytic DH domain (**Fig. 3C**). All three inhibitors showed comparable inhibition of Vav1 and Vav3, consistent with the high sequence homology of the two GEFs (**Fig. 3C,D and Supplementary Fig. 6,7,8,9**). Remarkably, only minimal inhibition was observed for the other GEFs tested, even for the closely related Vav2. Taken together, these results confirm the successful *de novo* design of effective and specific Vav1 inhibitors.

### Design of *de novo* inhibitors for Intersectin-2

We next extended the inhibitor design pipeline to Intersectin-2 (Itsn2). Unlike Vav1, there is no clearly defined auto-inhibitory domain for this GEF, and therefore, the only approach available was to use the RFDiffusion-ProteinMPNN-Alphafold2 pipeline to design *de novo* binders. As with the Vav1 pipeline, hotspots were used to guide interactions with both conserved and variable regions of the DH domain. Nine of ten Itsn2 designs expressed recombinantly (**Supplementary Table 3**), of which two exhibited measurable inhibition when tested at a high concentration of 100 µM using the nucleotide exchange assay with Itsn2 and Cdc42 (**Fig. 4A,B and Supplementary Fig. 10A**). Titration of the Itsn2_i2 design showed a dose response of inhibition with an IC_50_ of 23.8 μM (**Fig. 4C and Supplementary Fig. 10B**). Mutations introduced at the predicted binding interface of Itsn2_i2 abolished inhibitory activity, confirming that both inhibition and binding depend on the designed interaction surface (**Fig. 4 B,D,E and Supplementary Fig. 10C,D**). Finally, Itsn2_i2 showed no inhibition of Vav1 and GEFH1 exchange activity, confirming specificity towards Intersectin-2 (**Fig. 4F**).

**Figure 4.**
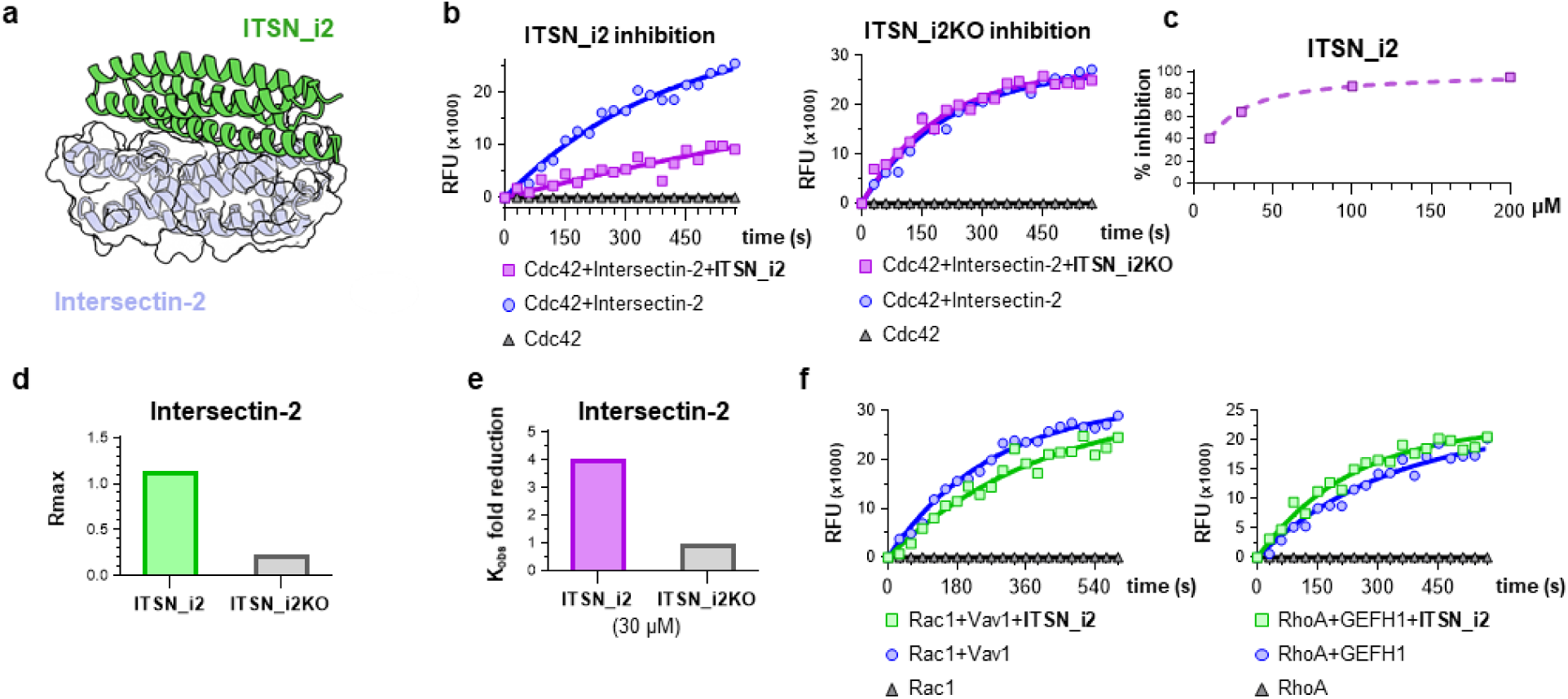
Design and validation of an engineered Intersectin-2 inhibitor. **a.** Model of engineered Intersectin-2 inhibitor Itsn_i2 (*green*) bound to the DH domain of Intersectin-2 (*cyan*). **b.** Inhibition of Intersectin-2 GEF activity towards Cdc42 by Itsn_i2 (*left*) and a knock-out version lacking the engineered contacts (*right*). **c.** Concentration dependent inhibition of Intersectin-2 by Itsn_i2 inhibitor. **d.** BLI binding of Itsn_i2 and Itsn_i2KO to recombinant Intersectin-2 DH-PH domain. **e.** Reduction in Intersectin-2/Cdc42 activity by Itsn_i2 and Itsn_i2KO (30 µM), as measured by changes in k_obs_ determined from **b**. **f.** Inhibition of Vav1/Rac1 and GEF-H1/RhoA by Itsn_i2.

### GEF inhibition in living cells

To quantify Vav1 inhibition in living cells, we employed a high-throughput, cell-based FRET biosensor assay that reports Rac1 activity in real time ^19^(**Fig. 5A,B**). The biosensor is based on fluorescence resonance energy transfer (FRET) between fluorophores attached to Rac1 and an associated Rac1-binding (AR) domain, which selectively interacts with Rac1 in its GTP-bound, active conformation. Upon GDP–GTP exchange catalyzed by guanine nucleotide exchange factors, Rac1 undergoes a conformational change that enables binding to the AR domain, resulting in increased FRET signal. Rac1 activity was quantified as the ratio of FRET to donor fluorescence (FRET/donor), providing a normalized measure of Rac1 activation. Vav1 was expressed as an emiRFP670-tagged construct to drive Rac1 activation in this cellular assay. Cells were co-transfected with different emiRFP670-coupled Vav1 variants, including wild-type full-length Vav1, a full-length construct in which regulatory tyrosines were mutated to glutamate, and two constructs containing only the minimal DH–PH-C1 domains. Among these, the full-length Vav1 mutant with tyrosine-to-glutamate substitutions produced the most robust Rac1 activation and was therefore selected for subsequent experiments (Fig. 5C). Cells were then co-transfected with this Vav1 construct and increasing DNA concentrations of three candidate inhibitors (Vav1_i6, Vav1_i50, and Vav1_i99) tagged with mScarlet3. Rac1 activity was quantified by measuring FRET efficiency (**Supplementary Fig. 11A,B**). Two control conditions, one in which Rac1 signaling was not inhibited and was used to define 100% Rac1 activity, and another where no GEF was transfected and was used to define 0% Rac activity, were used to normalize responses across experiments. Expression of Vav1_i6 and Vav1_i50 resulted in a dose-dependent reduction of Rac1 activity, as evidenced by a progressive decrease in FRET signal with increasing inhibitor DNA concentrations (**Fig. 5D**). This inhibitory effect was reproducible across replicates and consistent with effective suppression of Vav1-mediated Rac1 activation. In contrast, Vav1_i99 showed no significant effect on Rac1 activity across the tested concentration range, indicating a lack of inhibitory activity in this assay. To assess whether the observed effects were dependent on the inhibitors, we next performed the assay in cells using the respective knockout versions of Vav1_i6 and Vav1_i50 (**Fig. 5D**). Under these conditions, neither knockout altered Rac1 activity. These findings demonstrate that Vav1_i6 and Vav1_i50 selectively inhibit Vav1-dependent Rac1 activation.

**Figure 5.**
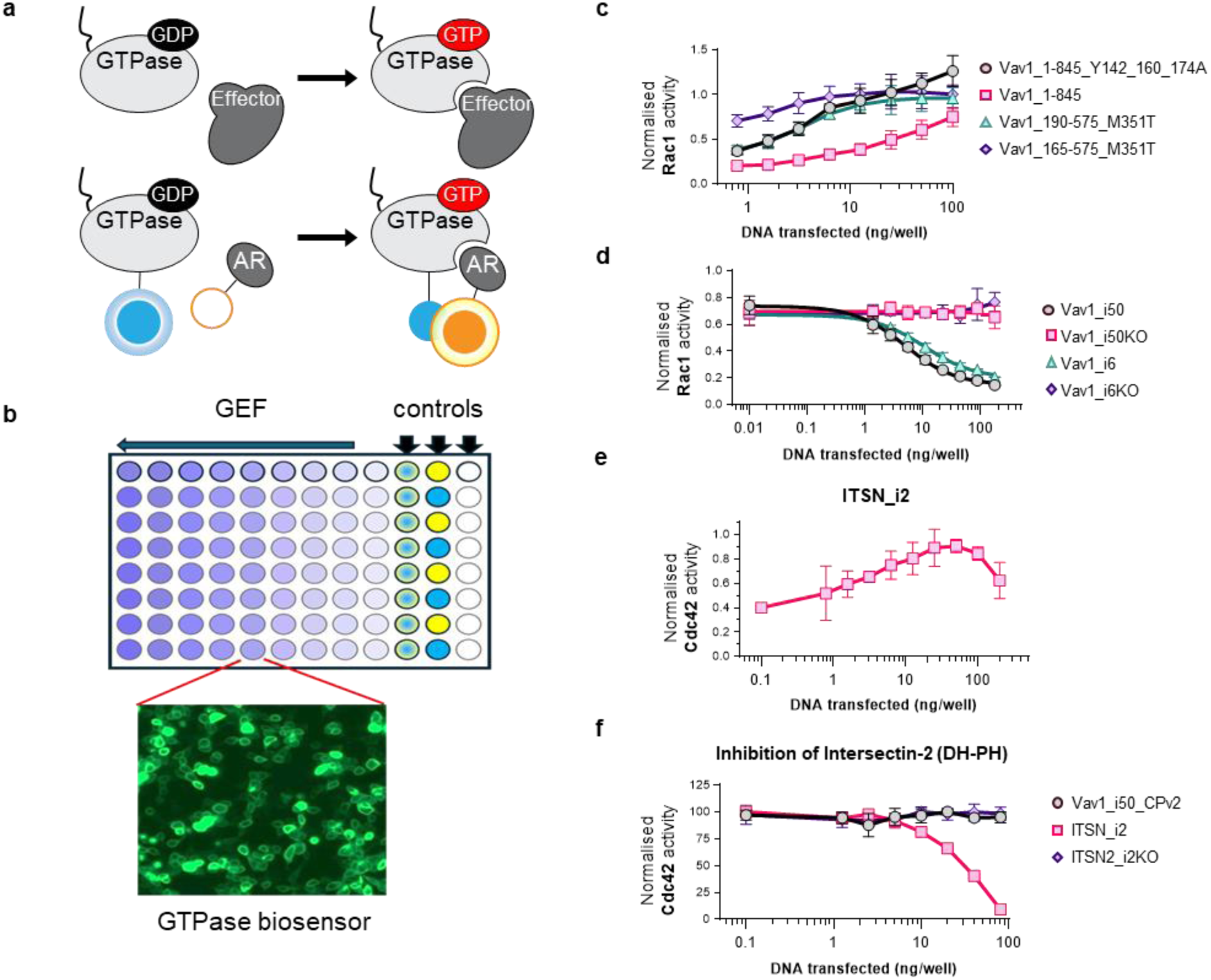
Inhibiton of GEF activity by engineered inhibitors in living cells. **a.** Schematic of the FRET-based biosensor used to quantify GTPase activation in living cells. **b.** Illustration of the high content 96-well plate assay used to measure GEF activity at different concentrations towards GTPase activation. Inset shows 293t cells transfected with Rac1 biosensors, Vav1, and inhibitor. **c.** Activation of Rac1 by constructs containing different Vav1 subdomains and mutations. **d.** Inhibition of Vav1 in living cells by inhibitors Vav1_i50, Vav1_i6, and their respective knock-out variants as reported by the Rac1 biosensor. **e.** Same plate assay as in **b**, showing Cdc42 FRET biosensor activity in cells upon expression of Intersectin-2. **f.** Inhibition of Intersectin-2 by Itsn_i2 inhibitor, its knock-out variant, and Vav1 inhibitor Vav1_i50, variant CPv2, as reported by the Cdc42 biosensor.

We next tested the ability of the intersectin inhibitor Itsn2_i2 to inhibit the GEF in living cells. Cells were transfected with various concentrations of Itsn2 together with the Cdc42 FRET biosensor, resulting in a dose dependent activation of Cdc42 (Fig. 5E)^20^. Next, Itsn2_i2 was transfected at various concentrations together with Intersectin-1 (Itsn1) (**Supplementary Fig. 11C**) or Intersectin-2 (Itsn2) (**Fig. 5F**). A knockout (KO) version of ITSN2_i2 was included as control. Itsn2_i2 exhibited a dose-dependent inhibition of Cdc42 activity in the presence of both Itsn1 and Itsn2, as measured by reduced FRET efficiency relative to control conditions, although inhibition was consistently more potent in the Itsn2-driven condition compared with Itsn1, as reflected by a steeper dose–response relationship. Lastly, we assessed whether Itsn2_i2 affects Vav1 in a Rac1 activation assay (**Supplementary Fig. 11B**). Itsn_i2 did not have any effect on Vav1 activity, supporting its specificity to Intersectin-2. This preferential activity toward Itsn2 is consistent with the structure-based design strategy, as Itsn2_i2 was computationally designed against Itsn2 rather than Itsn1. Using knockout control inhibitor, Itsn2_i2-KO had no detectable effect on Cdc42 activity, confirming that the observed inhibition is dependent on Itsn2-mediated Cdc42 activation.

We next sought to generate optogenetic probes from our Vav1 inhibitors by coupling them to light-activated modules that control their activity within cells. We used the Zlock design^21^, a previously described module composed of a Light-oxygen-voltage-sensing (LOV) domain and an engineered binder (Zdk2)^22^ that only interacts with the LOV domain only in the dark (**Fig. 6**), which can be used to control protein activity with light. For coupling of this optogenetic module to a GEF inhibitor the protein termini, where these modules are attached, need to flank the active site of the protein, i.e. the GEF binding interface of the inhibitor. To enable this strategy, we re-engineered Vav1_i50. The inhibitor was truncated, removing the C-terminal helix and used ProteinMPNN to redesign the newly exposed surfaces to aid folding and solubility, resulting in constructs Vav1_i50-DC (**Fig. 6A**). We then introduced a circular permutation, switching the 3rd and 5th helices so that the C-terminus of the inhibitor was now adjacent to the inhibitor/GEF interface (construct Vav1_i50-CP, **Fig. 6A**). This new re-engineered inhibitor was able to inhibit Vav1 in cells to the same extent as the original Vav1_i50 (**Fig. 6B**).

**Figure 6.**
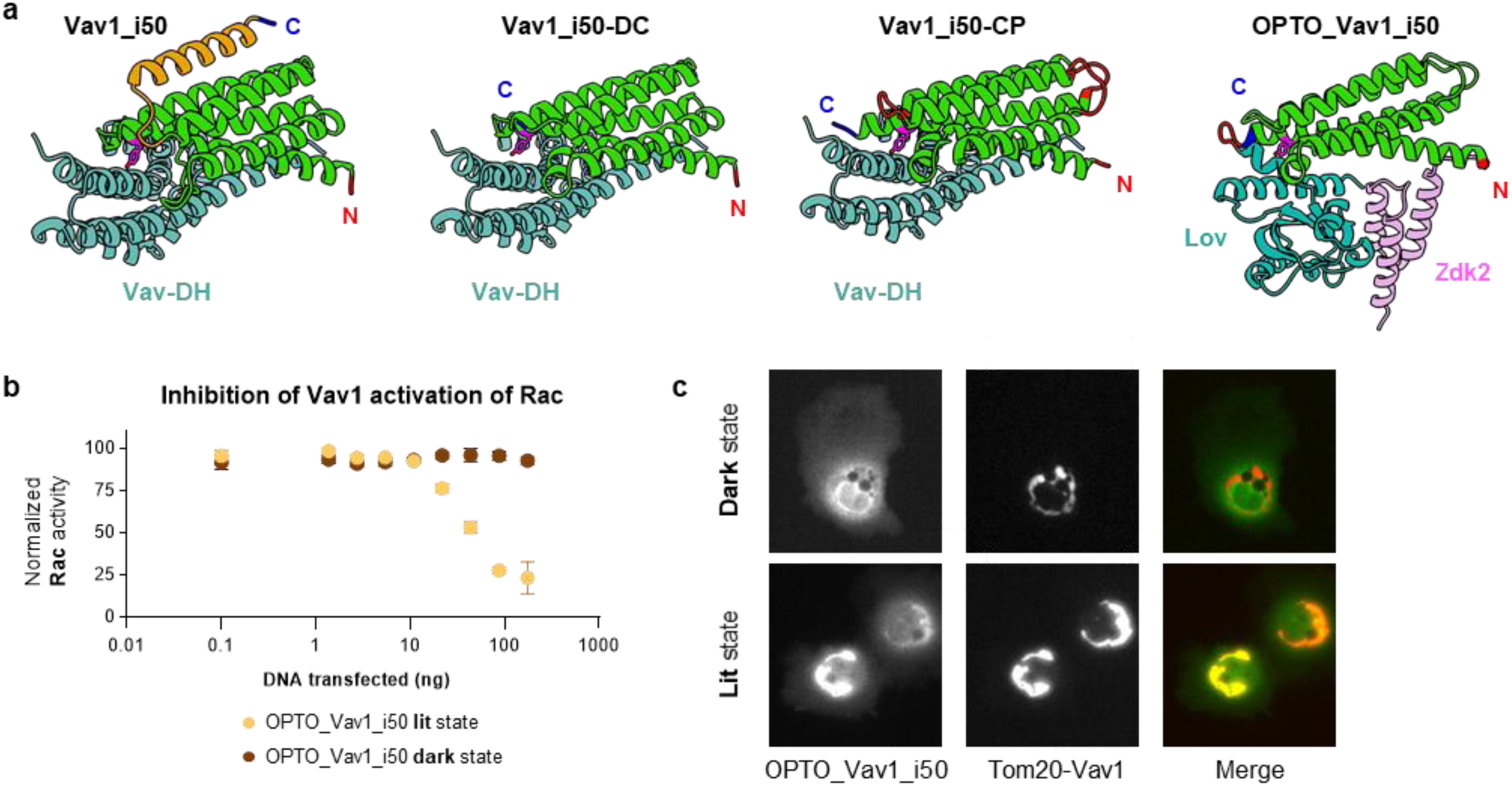
Functionalization of Vav1 inhibitor Vav1_i50 for optogenetic control. **a.** Redesign stages of Vav1_i50 structure for coupling to the LOVTrap optogenetic module. **b.** Inhibition of Vav1 activation of Rac1 in cells by OPTO_Vav1_i50 in its dark and irradiated states, simulated by mutations. **c.** Images of cells co-expressing OPTO_Vav1_i50 and Vav1 localized to the mitochondria. Vav1 (*center*) recruits OPTO_Vav1_i50 (*left*) to the mitochondria in its lit, but not dark state.

The Zlock components were then added to Vav1_i50-CP, with the Zdk2 at the N-termini and the LOV domain at the C-termini. We screened different linker lengths to optimize the difference between lit and dark state LOV mutants, so that the LOV domain interacted with the Zdk2 and inhibited Vav1 in cells only in the lit state but not in the dark state to produce the final OPTO_Vav1_i50 (**Fig. 6B**). We further validated this design by co-expressing OPTO_Vav1_i50 with a Vav3 construct that we had localized to the mitochondria with a Tom20 tag and showed that OPTO_Vav1_i50 only binds to the GEF in the lit state and not the dark state (**Fig. 6C**)

## Discussion

In this study, we established a generalizable strategy for engineering selective protein inhibitors of Dbl family guanine nucleotide exchange factors (GEFs), a class of signaling proteins that has been difficult to target. Through the combination of generative backbone design with structure-guided sequence optimization and validation, we generated novel protein inhibitors that engage the conserved DH domains of multiple GEFs and inhibit their catalytic activity. Our results demonstrate that protein design approaches can overcome challenges associated with targeting flat, highly conserved protein-protein interaction surfaces by small molecules, enabling both specificity and functional inhibition.

The catalytic DH domains of GEFs interact with small GTPases through extended surfaces that lack deep binding pockets typically exploited by small molecules. Moreover, the high degree of structural conservation across family members complicates efforts to achieve selectivity. These challenges are exemplified by the scarcity of reported inhibitors and by the limited specificity of existing small-molecule compounds. Our work shows that *de novo* designed protein inhibitors can circumvent these constraints by forming structures that compete with native regulatory interactions, providing a route to selective inhibition even within highly conserved protein families.

We pursued three complementary inhibitor design strategies. In the first approach, the Vav1 AID was grafted onto proteins of known structure from the Protein Databank. In the second method, we leveraged the known autoinhibitory domain (AID) of Vav1 as a starting motif and built proteins around it using RFDiffusion. ProteinMPNN was used to design sequences, and AlphaFold-Multimer was used to validate the conformation of the resulting proteins. This allowed the creation of additional *de novo* interactions to enhance affinity and specificity toward the DH domain. This approach effectively exploits endogenous regulatory motifs while extending them beyond their native context. In the third strategy, we designed inhibitors entirely *de novo* for Vav1, and Intersectin-2 without relying on natural templates. Although less effective, this method proved capable of generating stable binders that inhibit GEF activity, demonstrating that functional inhibitors can be engineered even in the absence of detailed knowledge of native regulation.

Importantly, Vav1 inhibitors showed specificity in biochemical assays, selectively inhibiting their intended targets and not reacting with related GEFs. This specificity likely arises from the use of hotspot residues as guides for the inhibitor design. *De novo* inhibitors designed against Intersectin-2 had lower affinity. Nevertheless one Intersectin-2 inhibitor showed 50% inhibition of the GEF *in vitro* and complete inhibition in cells. Therefore, inhibition is maintained in the complex molecular environment of living cells, supporting the idea that these designed proteins are sufficiently stable, specific, and functional to operate *in vivo*.

In order to have control over the spatio-temporal kinetics of GEFs, we decided to functionalize the inhibitors with optogenetic probes for live-cell imaging. Optogenetic probes rely on the use of modules that can be activated under light and turned off under dark conditions. One of the Vav1 inhibitors, Vav1_i50, was best fitted structurally to accommodate the Zlock and Light-oxygen-voltage-sensing domains, and was thus developed into an optogenetic tool. The re-engineered inhibitor was able to inhibit Vav1 in cells to the same extent as the parental protein, proving that the modifications to accommodate the domains did not impact its functional activity. This tool will allow the assessment of the morphological and signaling changes upon Vav1 inhibition in real-time, under the control of the user, who can turn on/off the inhibitors in discrete cell regions and at any chosen times during the experiment. Together, these findings suggest that generative protein design can move beyond producing binders and yield biologically active modulators of intracellular signaling pathways.

Our work establishes a versatile framework for targeting Dbl family GEFs and, more broadly, other signaling proteins dominated by flat protein–protein interaction surfaces. By integrating generative backbone design, sequence optimization, and functional validation, this platform enables the rational creation of inhibitors that were previously inaccessible using traditional approaches. Beyond GEF biology, this strategy should be broadly applicable to dissecting signaling networks, probing the roles of individual family members in complex cellular contexts, and developing new classes of biologically inspired inhibitors for challenging protein targets.

## Methods

### Design of inhibitors by side chain grafting

A database of 9884 candidate scaffolds was created by selecting from the Protein Data Bank (PDB) structures that were: 1) determined by x-ray crystallography; 2) high resolution (<2.8 Å); 3) monomeric; 4) expressed in *E. coli*; 5) not of human origin and 6) that did not contain ligands. Epitope scaffolds were designed with Rosetta using the RosettaScripts for side chain grafting described in *Silva et. al*., with the following parameters in the *MotifGraft* mover: RMSD_tolerance = “0.5”; NC_points_RMSD_tolerance = “0.5”; clash_test_residue = “ALA”; clash_score_cutoff = “5”, hotspots=“1:2:5:6”. For the design of auto-inhibitory domain (AID) epitope scaffolds, the structure of AID^173–178^ fragment ^173^IYEDLM^178^ was isolated from the crystal structure of the Autoinhibited-Vav1 (PDBid: 3KY9), keeping the identity of all the epitope residues the same. From the scaffold generated by the automated protocol, the top 100 models by Rosetta ddG that were also smaller than 150 amino acids were visually examined to ensure appropriate motif transplantation. Additional changes were introduced in the inhibitor scaffold using Rosetta fixed backbone design to remove Vav1-inhibitor scaffold contacts that were not due to the binding interface. The best 7 designs were chosen for experimental characterization.

### Design of inhibitors using de-novo design

A fully automated, modular computational pipeline was developed to integrate backbone generation, sequence design, and structure prediction into a single workflow. The pipeline consisted of three main stages: (i) backbone generation using RFdiffusion, (ii) sequence optimization with ProteinMPNN, and (iii) structure prediction using AlphaFold Multimer, followed by a unified analysis and ranking module.

To make Vav1 AID-templated inhibitors, RFdiffusion was run in conditional mode using PDB:3KY9 with the contig specification [40–100 / A169–178 / 40–100 / 0 B192–390], which held residues A169–A178 (AID sequence EGDEIYEDLM) fixed between two de novo design segments of 40–100 residues each, and kept B192–B390 (the Vav1 DH domain) fixed as the target protein. Interface guidance was provided via hotspot specification (ppi.hotspot_res = [B201, B313, B319, B320, B331]) to bias backbone sampling toward productive contacts with DH. Each RFdiffusion job generated 100 candidate backbones (inference.num_designs = 100), which were then passed forward as the structural templates for sequence design.

Analogous setups were applied to de novo designs targeting Vav1, GEFH1, and Intersectin-2 in the absence of the AID by adjusting receptor-specific hotspot definitions, enabling direct reuse of the same pipeline logic across multiple targets. The following parameters were used:

- GEF-H1 DH domain (PDB:7G80): “[B204-448/0 150-150]” and hotspots “[B376,B378,B385,B399]”, 15 designs

- Intersectin-2 (PDB:3GF9): “[A1178-1367/0 150-150]” and hotspots “[A1306,A1313,A1320,A1328]”, 15 designs

- Vav1 (PDB:3KY9): “[B192-508/0 150-150]” and hotspots “[B212,B320,B327,B331]”, 10 designs

ProteinMPNN was used to optimize amino-acid sequences for each backbone while keeping the AID residues fixed. Automated FASTA processing ensured a one-to-one correspondence between RFdiffusion backbones, ProteinMPNN sequences, and AlphaFold inputs, eliminating manual intervention and preventing sequence–structure mismatches.Designed sequences were evaluated using AlphaFold Multimer. Predicted structures were analyzed using a custom Python framework that computed structural confidence (mean pLDDT), agreement with the designed backbone (RMSD and TM-score), interfacial confidence (ipTM/pTM, PAE), steric clashes, and topological descriptors. Designs were ranked using a composite score emphasizing both local confidence and global structural agreement:

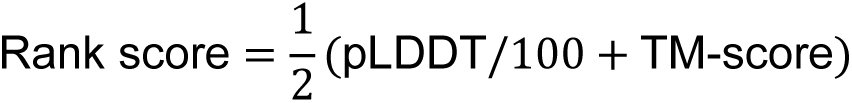

All stages were executed on a high-performance computing cluster, enabling rapid, high-throughput design. The same pipeline was applied without modification across multiple targets, yielding ∼20 high-confidence designs per target after automated filtering.

### Inhibitors and GTPases expression and purification

Plasmids encoding His-tagged inhibitors and GTPases were transformed into *E. coli* BL-21 (DE3) (New England Biolabs) cells and 5 mL starter cultures were grown overnight at 37°C in Lysogeny Broth (LB) supplemented with 50 μg/mL kanamycin. The cultures were diluted 1:100 in Terrific Broth TB media (RPI) supplemented with kanamycin and grown at 37°C to an OD600 of ∼0.6. The temperature was subsequently lowered to 16°C and the cultures were grown to OD600 of ∼0.8 and induced with isopropyl β-D-1-thiogalactopyranoside (IPTG) to a final concentration of 0.5 mM. The cultures were shaken for 16–18 hours at 200 rpm. Cell pellets were collected by centrifugation at 14,500 x g and lysed in Lysis buffer (B-PER Reagent (ThermoFisher Scientific), 5 mM MgCl2, 2 mM TCEP, 1 mM PMSF plus protease inhibitor). The lysates were centrifuged at 14,000 x g for 30 minutes at 4°C. The supernatant was incubated with Ni-NTA beads (Qiagen) equilibrated with lysis buffer for an hour at 4°C. The beads were settled by centrifugation, and the supernatant was removed by pipetting. The beads were washed with wash buffer (20 mM Tris-HCl, 150 mM NaCl, 5 mM MgCl_2_, 2 mM TCEP, 30 mM imidazole, 5% glycerol, pH=7.5) after which the protein was eluted with elution buffer (20 mM Tris-HCl, 150 mM NaCl, 5 mM MgCl_2_, 2 mM TCEP, 250 mM imidazole, 5% glycerol, pH=7.5). Protein expression and purity was confirmed by SDS-PAGE analysis, and quantified spectro-photochemically at 280 nm on a Nanodrop 2000 (ThermoFisher Scientific). The eluted protein was concentrated on 3kDa MW spinMW spin columns (ThermoFisher Scientific) and buffer exchanged in Exchange Buffer (20 mM Tris-HCl,150 mM NaCl,1 mM MgCl_2_,1 mM TCEP, 5% glycerol, pH=7.5). For further biochemical characterization, inhibitors were purified by size exclusion chromatography in 20 mM Tris-HCl,150 mM NaCl,1 mM MgCl_2_,1 mM TCEP, 5% glycerol, pH=7.5, on an AKTA-Go FPLC (Cytiva) using a Superdex 200 Increase 10/300GL column. Fractions containing monomeric protein were pooled and concentrated as above.

### Guanine-Exchange Factors expression and purification

Plasmids encoding His-tagged GEFs were transformed into *E. coli* BL-21 (DE3) (New England Biolabs) cells and 5 mL starter cultures were grown overnight at 37°C in Lysogeny Broth (LB) supplemented with 50 μg/mL kanamycin. The cultures were diluted 1:100 in Terrific Broth TB media (RPI) supplemented with kanamycin and grown at 37°C to an OD600 of ∼0.6. The temperature was subsequently lowered to 16°C (22°C for Vav1, Vav2, Vav3) and the cultures were grown to OD600 of ∼0.8 and induced with isopropyl β-D-1-thiogalactopyranoside (IPTG) to a final concentration of 0.5 mM. The cultures were shaken for 16–18 hours at 200 rpm. Cell pellets were collected by centrifugation at 14,500 x g and lysed in Lysis buffer (B-PER Reagent (ThermoFisher Scientific), 5 mM MgCl2, 2 mM TCEP, 1 mM PMSF plus protease inhibitor). The lysates were centrifuged at 14,000 x g for 30 minutes at 4°C. The supernatant was incubated with Ni-NTA beads (Qiagen) equilibrated with lysis buffer for an hour at 4°C. The beads were settled by centrifugation, and the supernatant was removed by pipetting. The beads were washed with wash buffer (50 mM HEPES,100 mM NaCl, 5 mM MgCl_2_, 30 mM imidazole, 5% glycerol, 2 mM TCEP, pH=7.5) after which the protein was eluted with elution buffer (50 mM HEPES, 100 mM NaCl, 5 mM MgCl_2_, 250 mM imidazole, 5% glycerol, 2 mM TCEP, pH=7.5). Protein expression and purity was confirmed by SDS-PAGE analysis, and quantified spectro-photochemically at 280 nm on a Nanodrop 2000 (ThermoFisher Scientific). The eluted protein was concentrated on 3kDa MW spinMW spin columns (ThermoFisher Scientific) and buffer exchanged in Exchange Buffer (50 mM HEPES, 100 mM NaCl, 1mM MgCl_2_, 5% glycerol, 1 mM TCEP).

### Differential Scanning Fluorimetry

Samples were analyzed using a Prometheus Panta (Nanotemper Technologies). All samples were diluted to 0.5 mg/mL and 15 µL was added to glass capillaries in triplicate for measurements. The instrument was programmed to measure 350 nm and 330 nm fluorescence from 15.0-110.0°C. All curves reported represent averages from the triplicate readings.

### BODIPY GDP exchange assay

Rac-1 and Vav1 were used at the concentrations of 1.6 µM and 0.2 µM, while RhoA, Cdc42 were run at 10 µM and GEFH1 and Intersectin-2 at 0.5 µM. Reaction mixtures contained the GTPase and 100 nM BODIPY FL GDP (Thermo Fisher Scientific) in 700 µl assay buffer (20 mM Tris, 150 mM NaCl, 2 mM DTT, 5 mM MgCl_2_, 5% glycerol and 0.01% NP-40, pH 7.5). Samples were read on a spectrofluorometer (FS5, Edinburgh Instruments) by recording emission at 511 nm upon 500 nm excitation. GEFs and inhibitors were pre-incubated at RT for 15 minutes. Upon baseline stabilization the GEF and inhibitor were added to the reaction mixture. Time-course data was plotted starting from 0s for Vav1 and GEFH1, while Intersectin-2 manifested a drop in signal upon the addition of the inhibitor, and was thus plotted starting from 90s. Time-course data was analyzed by non-linear regression using a one-phase association model. The model estimates an observed rate constant (K), which represents the apparent nucleotide-exchange rate, and the corresponding time constant (τ), defined as τ = 1/K. K directly reflects the kinetics of the reaction, providing a quantitative measure of inhibitor-induced slowing.Fits were performed globally with the initial signal (Y₀) constrained to zero and the plateau shared across conditions, while K was allowed to vary. Inhibitory effects were quantified as the percent reduction in K relative to the negative control, calculated as % inhibition = (1 − K_inhibitor_/K_control_)x100. The time constant τ was reported as a derived measure of the characteristic reaction timescale. This approach enabled direct comparison of reaction kinetics across conditions independent of differences in maximal signal.

### Biotinylation with Avi-tag

Biotinylation was performed using a BirA biotin-protein ligation kit (Avidity) on proteins produced with a C-terminal avidin tag sequence (GLNDIFEAQKIEWH) for RF_50_C61A_C133A and N-terminal avidin tag for RF_6 and RF_99_V79A_L112A_L114A. Following addition of enzyme kit reagents, proteins were incubated for 5h at 30°C. To remove excess biotin, proteins were transferred to 10kDa MW spin columns (Amicon) and five washes were performed in PBS 1X pH7.4 (Gibco). Final protein concentration was measured using NanoDrop.

### Bio-layer Interferometry (BLI)

To assess binding between inhibitors and Vav1, BLI was employed using Octet R4 (Sartorius). Biotinylated inhibitors were linked onto SA biosensors (Octer SA Biosensors, Sartorius, 18-5019) at a concentration of 20 μg/mL diluted in running buffer (20 mM Tris, 150 mM NaCl, 1% BSA, 0.05% Tween and 0.2 M Sucrose) with a loading time of 600 seconds. The sensors were exposed to various increasing concentrations of Vav1 for 300 seconds to allow association and 300 seconds for dissociation. GEFH1 inhibitors were run in similar conditions, at a concentration of 85 μg/mL in running buffer (1X HBS with 0.2% BSA). Data analysis Octet^®^ Analysis Studio software was used to analyze the results and measure kinetics and affinity, with reference sensor and zero-analyte control subtracted from the raw data.

### Cell based plate assay/Testing inhibitors using high-throughput microscopy

#### Image acquisition

Cells were imaged and analyzed as previously described^23^. HEK-293t cells were plated in 96-well plates with flat µ-clear plastic bottoms (Greiner bio-one) coated with poly-l-lysine (Sigma). DNA for biosensors for Rac1, Cdc42 or RhoA^24^, along with the minimum amount of emiRFP670 tagged GEF (Vav1, Intersectin-2, or Gef-H1) to maximally activate the biosensor, and with varying amounts of mScarlet3 tagged inhibitor were combined, and then transfection complexes were formed in 96-well plates prior to transfection. Cells were transfected in triplicate. After 24 hrs of expression, growth media was replaced with HBSS (Sigma) containing 1% FBS and 10mM HEPES (Gibco), prior to imaging. Cells were imaged using a Nikon Lambda D 10X, 0.45 NA objective on a Nikon Ti2 inverted microscope and using Elements acquisition software and Spectra III light engine illumination (Lumencor). Excitation filter: ZET405/445/514/561/640x (Chroma). Dichroic mirror: 89903bs (Chroma). Emission filters used (all Chroma) were CFP: 473/24, FRET/YFP: Em 540/24, mScarlet3: Em 595/33, emiRFP670: Em 697/60. Images were obtained on a Orca-Fusion sCMOS camera (Hamamatsu).

### Image analysis and data processing

Images were analyzed using MATLAB (Mathworks). Briefly, 3 fields were imaged for each well and the intensity was summed for each channel. These intensities were then background subtracted using values from wells that were mock transfected, and ratios were calculated using these background-subtracted values. For these experiments we plot R against DNA transfected: R= (FRET – α(Donor) – β(Acceptor))/Donor where R is the Ratio, FRET is the total FRET intensity as measured, α is the bleed-through of the donor into the FRET signal, β is the bleed-through of acceptor into the FRET signal, and Donor, Acceptor. Curves from different days were normalized such that values for no inhibitor are equal to 0% inhibition and wells with no GEF are 100% inhibition.

## Data analysis

Statistical analysis was performed with GraphPad Prism 8.

## Acknowledgements

The authors would like to thank A. Brenda Kapingidza for her help with protein expression, and Jared Lindenberger for assistance with the DSF experiments. We thank Silas Pontes de Almeida for graphical help with the structure panels in the figures.

## Authors contributions

M.L.A., F.V., and D.J.M H.B designed the inhibitors computationally. F.V., I.D., C.H., and J.C. expressed and purified the proteins recombinantly. F.V. and C.H. conducted the *in vitro* GEF activity assays. F.V. and J.C. carried out BLI measurements. F.V. conducted the DSF analysis.

D.W. performed SEC analyses. D.J.M. conducted the *in cellulo* live-cell inhibition assays. P.K. and H.B. developed and implemented an automated pipeline for *de novo* inhibitor design. F.V., D.J.M and M.L.A. wrote the first draft of the manuscript. F.V. and C.H. developed the figures. All authors reviewed and approved the final version of the manuscript. M.L.A. coordinated the project.

## Code availability statement

The protein design pipeline code was implemented in Python v3.9+ and is publicly available Github: https://github.com/HemanB/High-Throughput-Protein-Design

---

**Supplementary Figure 1.**
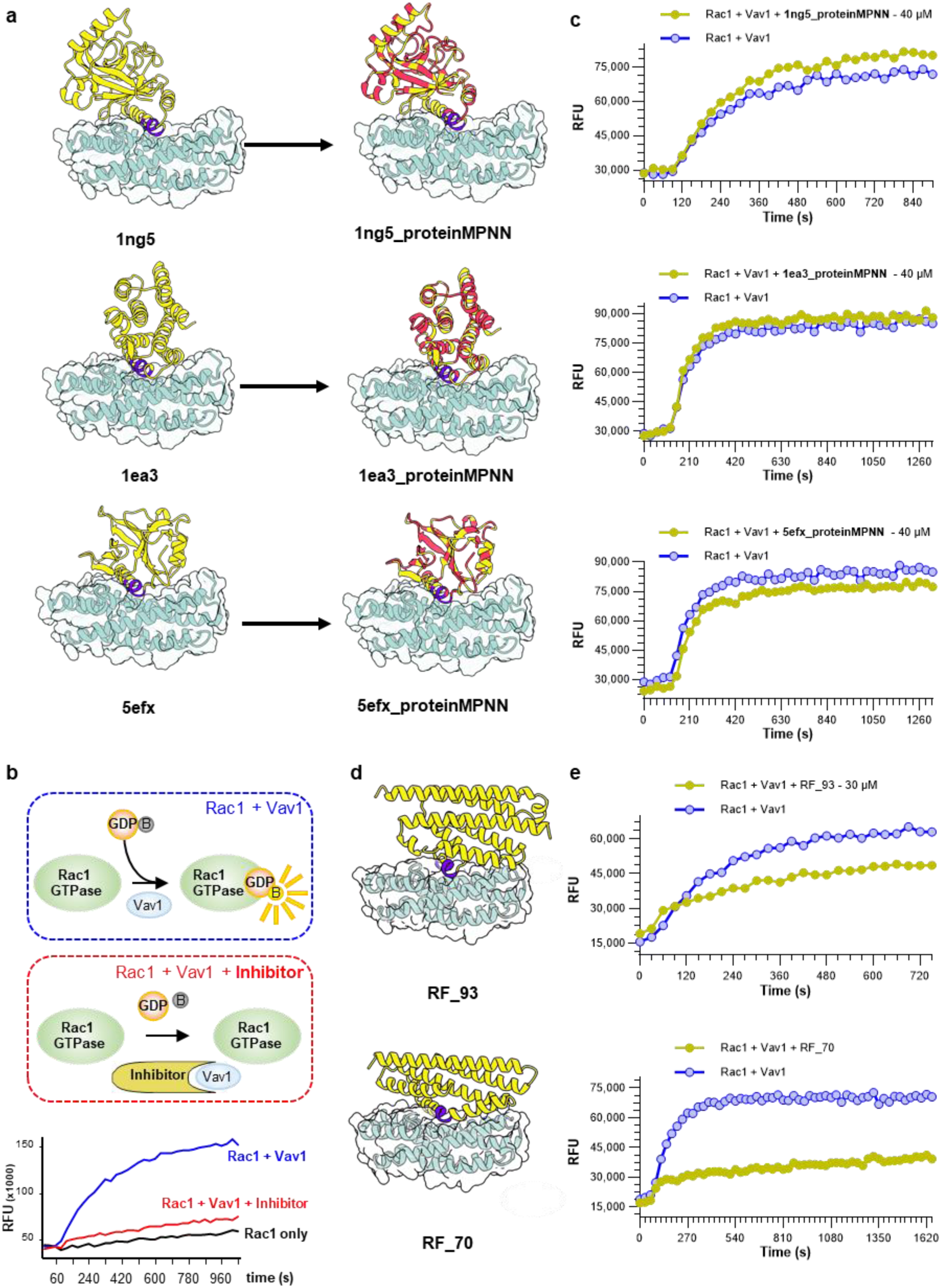
Models and characterization of Vav inhibitors developed by motif grafting. **a.** *Left*: inhibitor models (Vav1_inh_1ea3, 1ng5, and 5efx; *yellow*) displaying the autoinhibitory domain (AID, *purple*) bound to the Vav1 DH domain (*cyan*). *Right*: the same inhibitors after sequence optimization using ProteinMPNN. Amino acid substitutions introduced by ProteinMPNN relative to the original Rosetta designs are highlighted in red. **b.** Schematic of the BODIPY-GDP fluorescence-based nucleotide exchange assay with typical data. **c.** Vav1 activity towards Rac1 the absence (*blue*) and presence (*olive*) of the inhibitor designs 1ng5_proteinMPNN, 1ea3_proteinMPNN, 5efx_proteinMPNN (40 µM) from **a**. **d.** Models of Vav1 inhibitors Vav1_i93 and Vav1_i70 colored as in **a**. **e.** Partial inhibition of Vav1 by Vav1_i93 and Vav1_i70 at 30 µM (*olive*) compared to Vav1 alone (*blue*

**Supplementary Figure 2.**
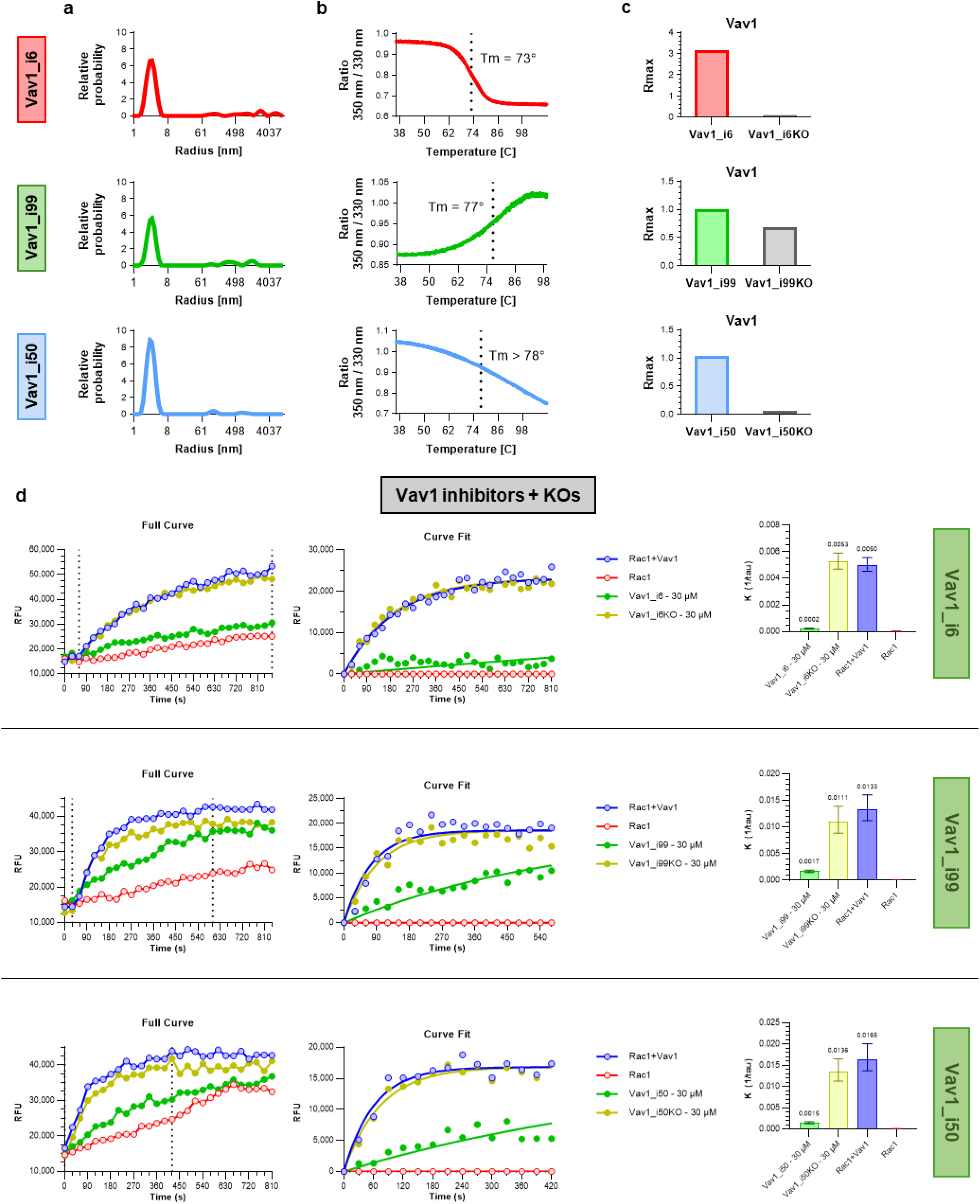
Biophysical characterization of Vav1 inhibitors. a. Dynamic light scattering (DLS) analysis showing a single dominant species with an average hydrodynamic radius of ∼3 nm. b. Differential scanning fluorimetry (DSF) measurements to measure the thermal stability of the inhibitor designs. c. Bio-layer interferometry (BLI) analysis showing R_max binding responses of each inhibitor and the corresponding knockout construct to Vav1. Inhibitors were tested at 10.13 µM. d. Vav1 inhibition by Vav1_i6 (*upper*), Vav1_i99 (*middle*), and Vav1_i50 (*lower*), tested at 30 µM. Each condition includes Rac1 + Vav1 + inhibitor (*green*), Rac1 + Vav1 + inhibitor knockout (*olive*), Rac1 + Vav1 positive control (*blue*), and Rac1 alone negative control (*red). Left*: collected data; *Center*: curve fittings. *Right*: k_obs_ values determined from **b**.

**Supplementary Figure 3.**
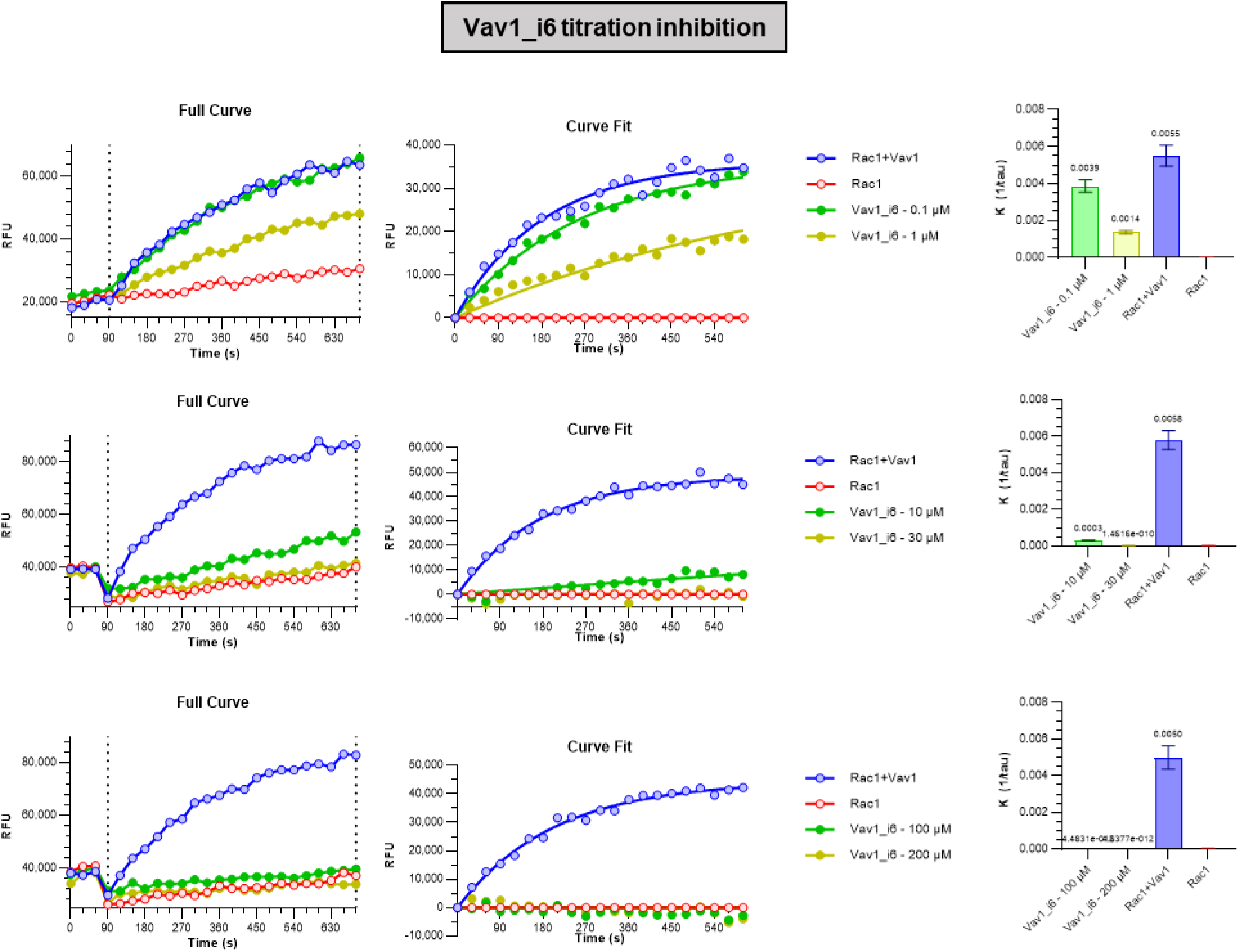
**Dose response of Vav1 inhibition by Vav1_i6.**

**Supplementary Figure 4.**
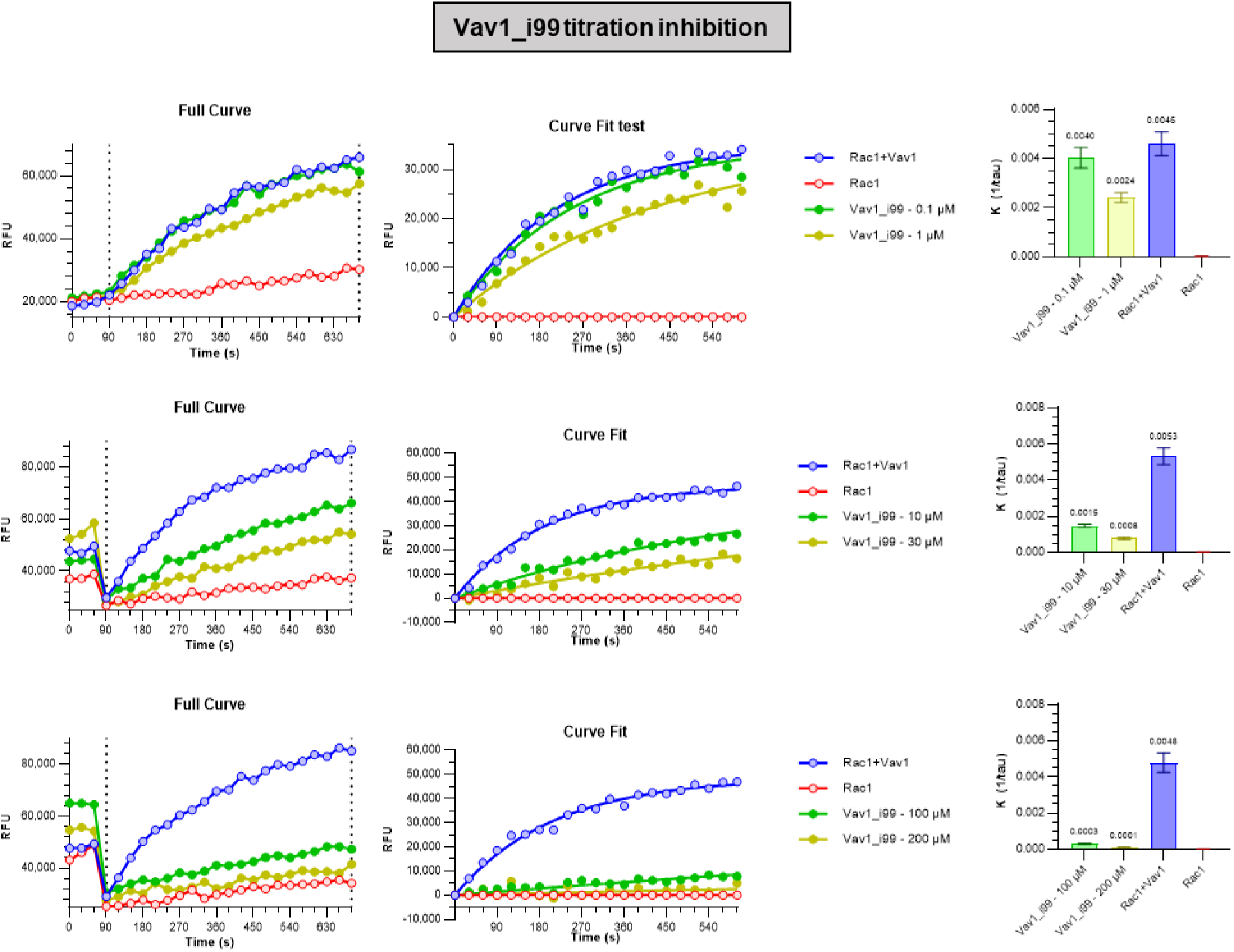
**Dose response of Vav1 inhibition by Vav1_i99.**

**Supplementary Figure 5.**
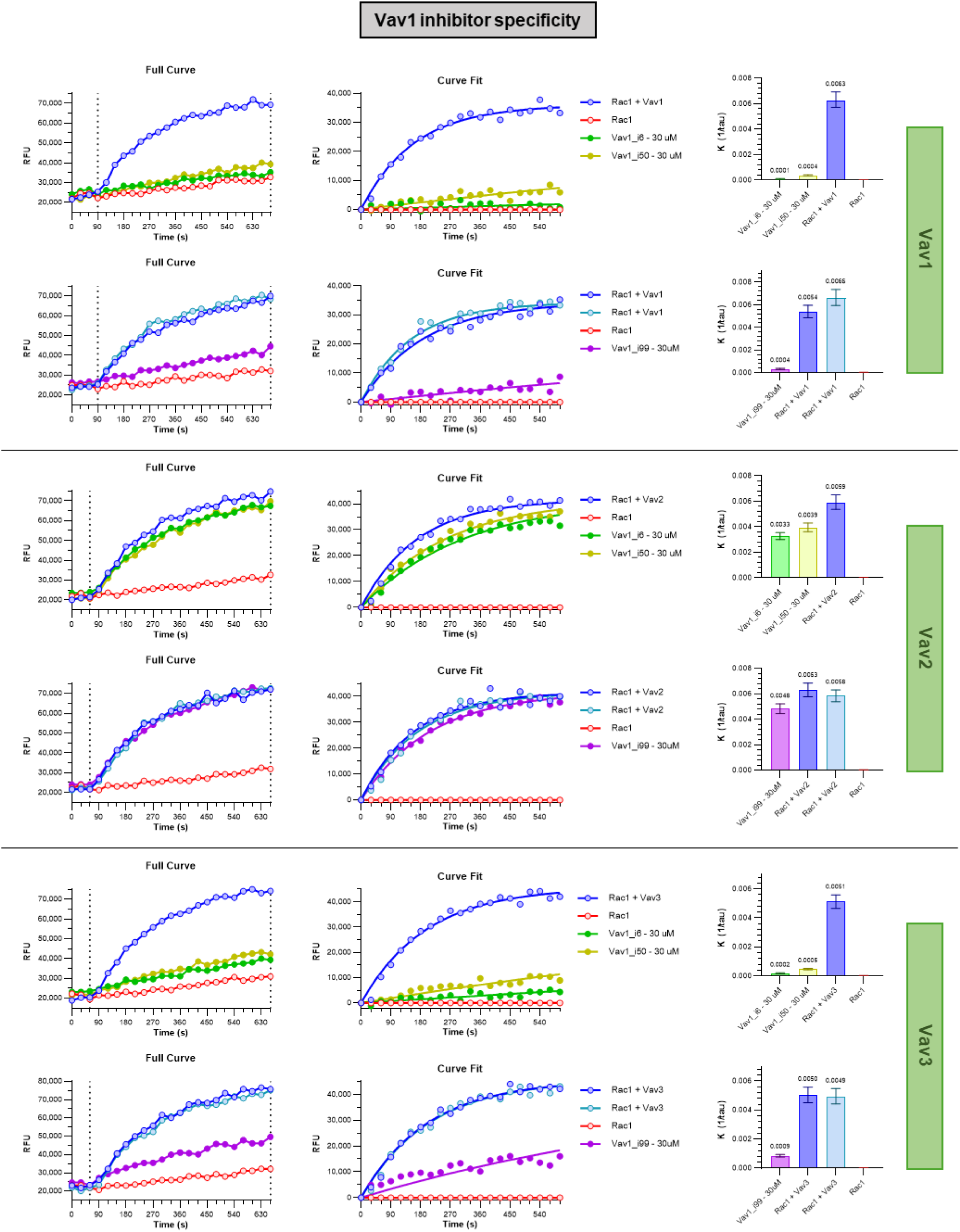
**Dose response of Vav1 inhibition by Vav1_i50.**

**Supplementary Figure 6.**
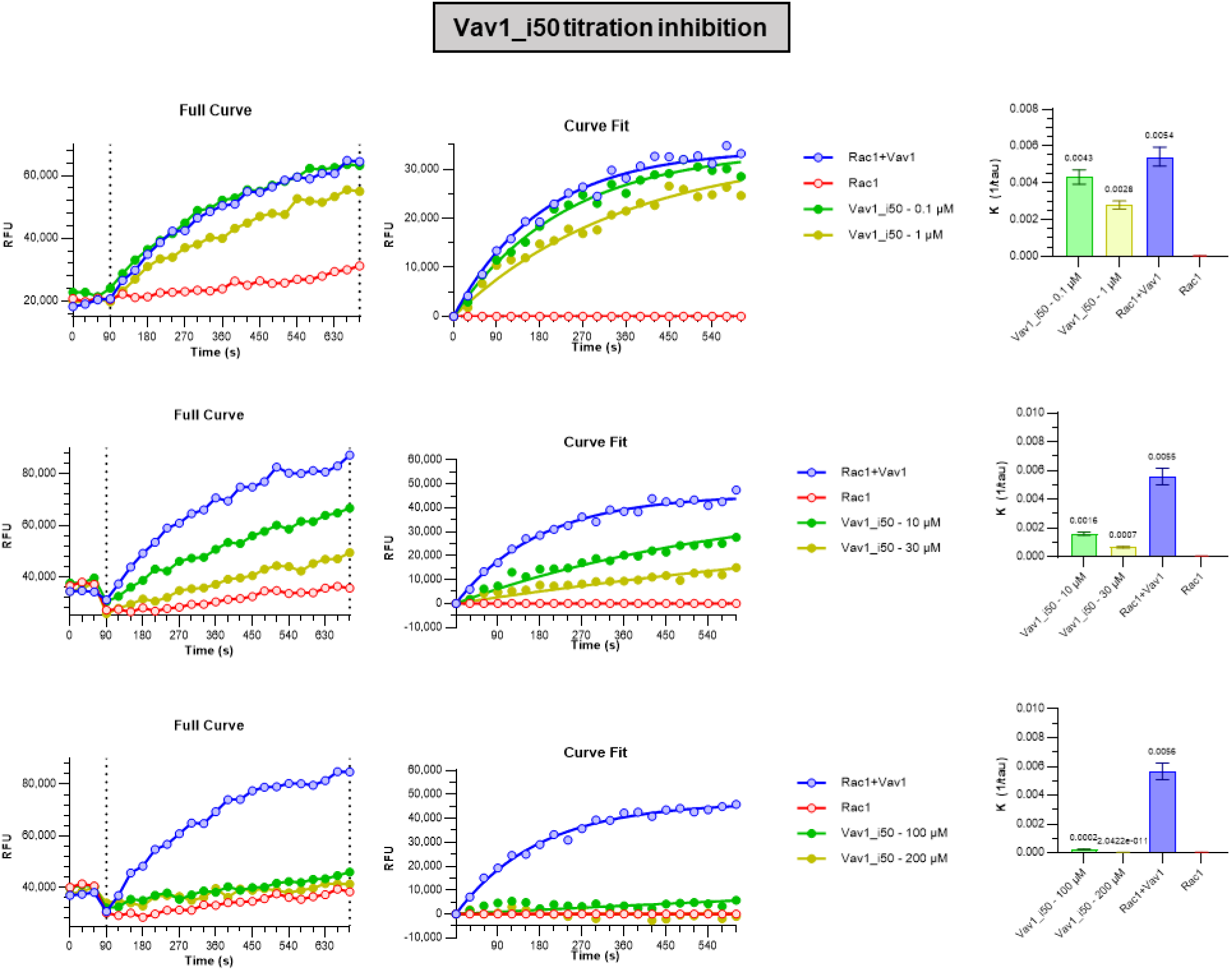
**Inhibition of different Dbl family GEFs by engineered Vav1 inhibitors.**

**Supplementary Figure 7.**
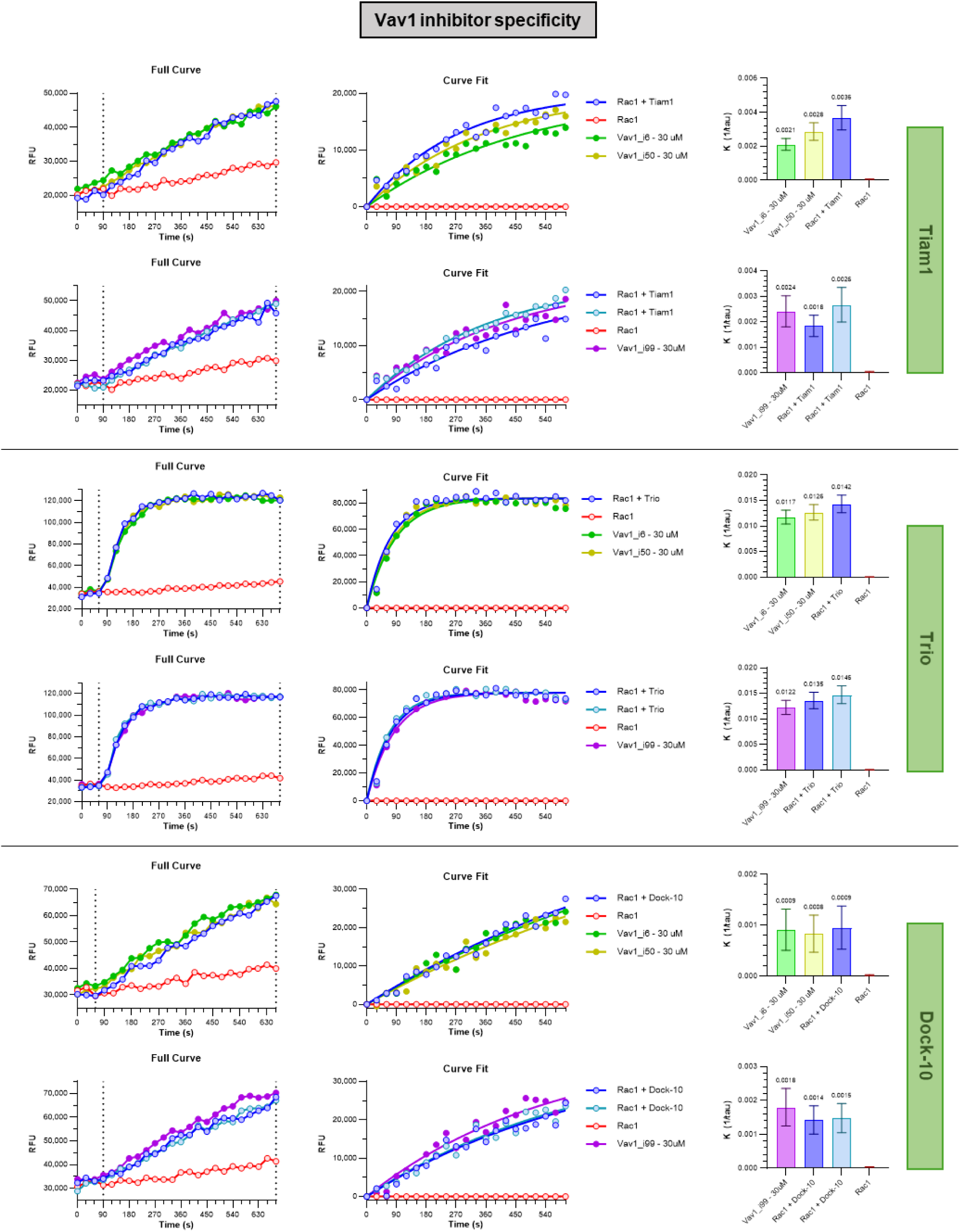
**Inhibition of different Dbl family GEFs by engineered Vav1 inhibitors.**

**Supplementary Figure 8.**
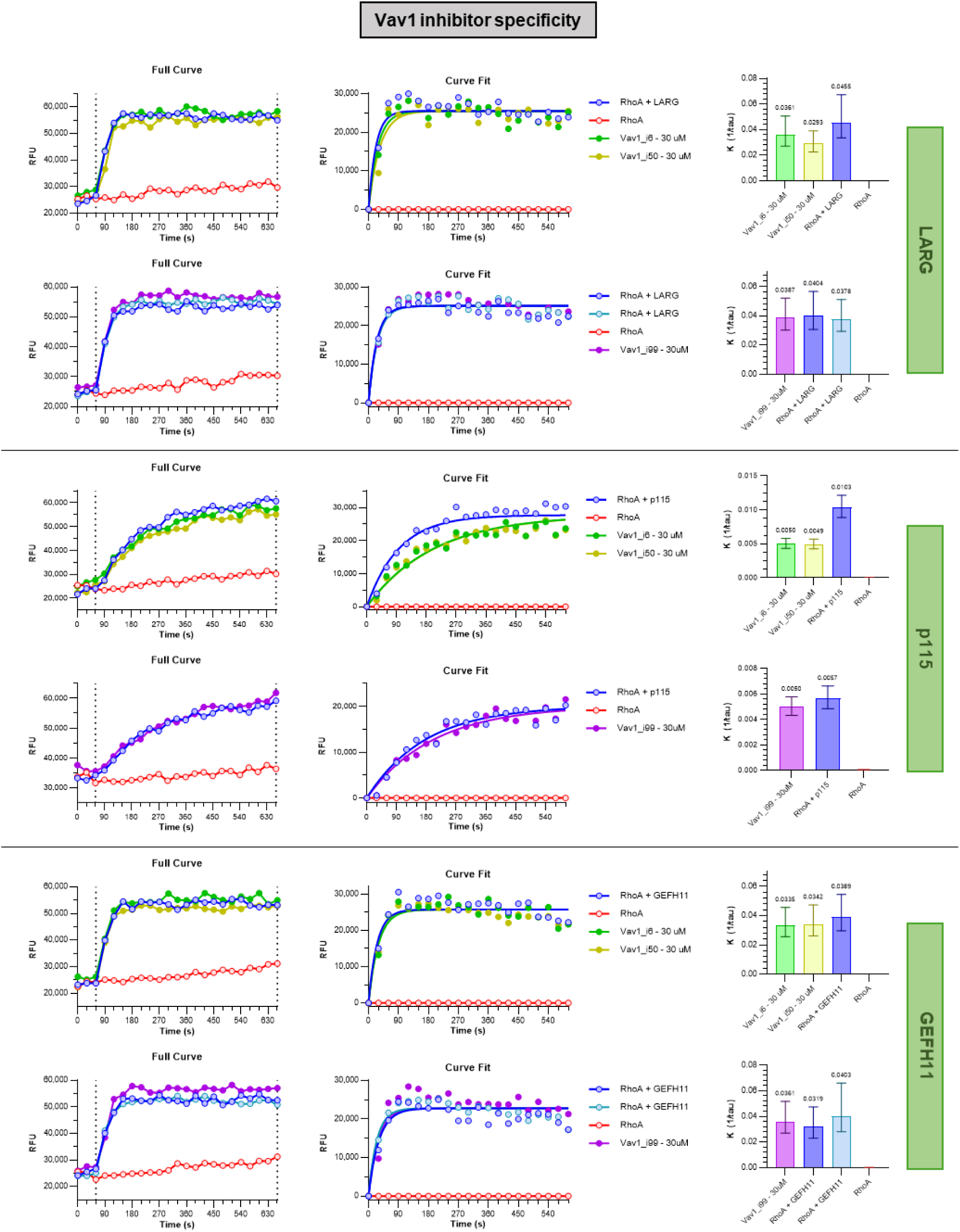
**Inhibition of different Dbl family GEFs by engineered Vav1 inhibitors.**

**Supplementary Figure 9.**
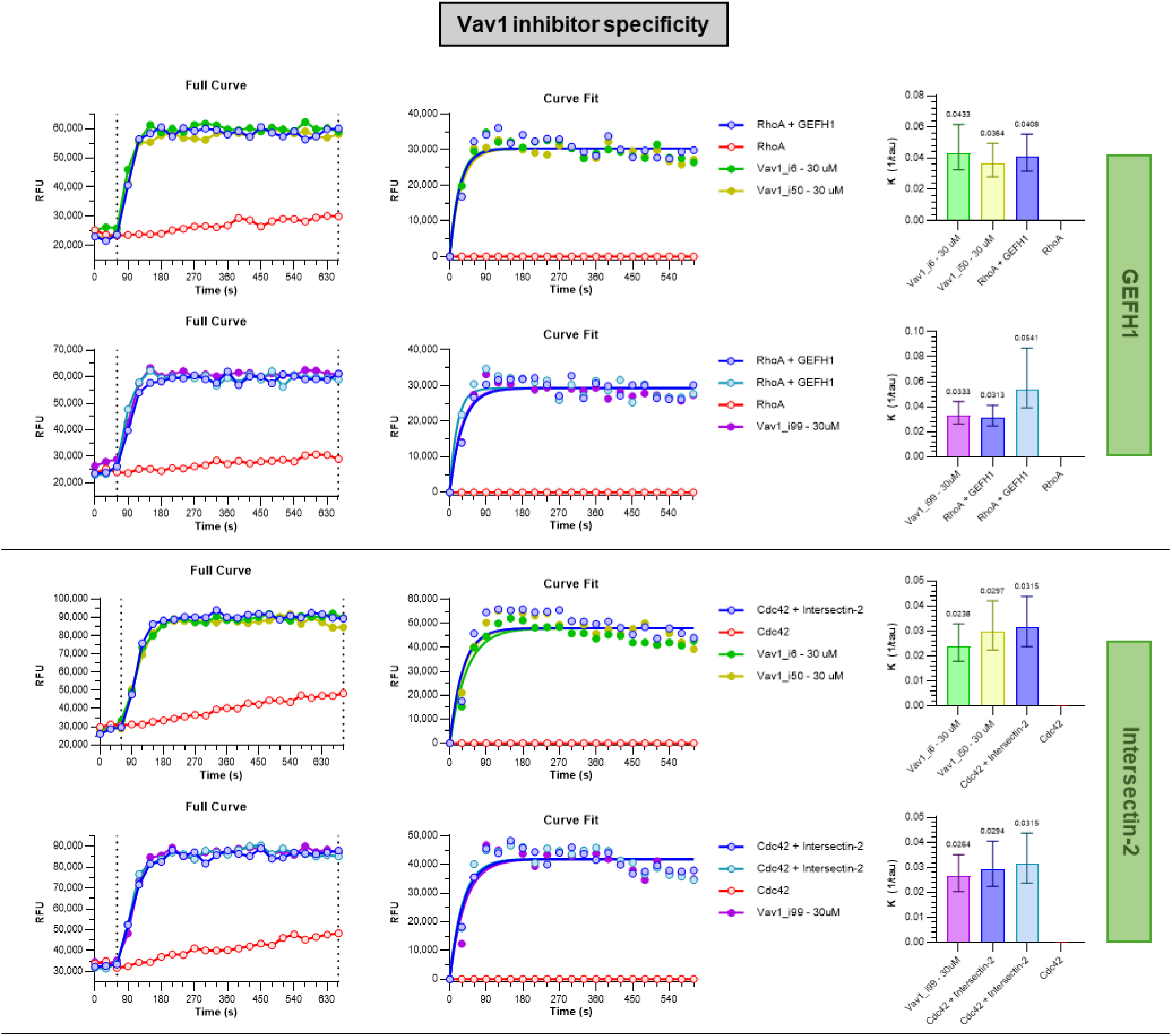
**Inhibition of different Dbl family GEFs by engineered Vav1 inhibitors.**

**Supplementary Figure 10.**
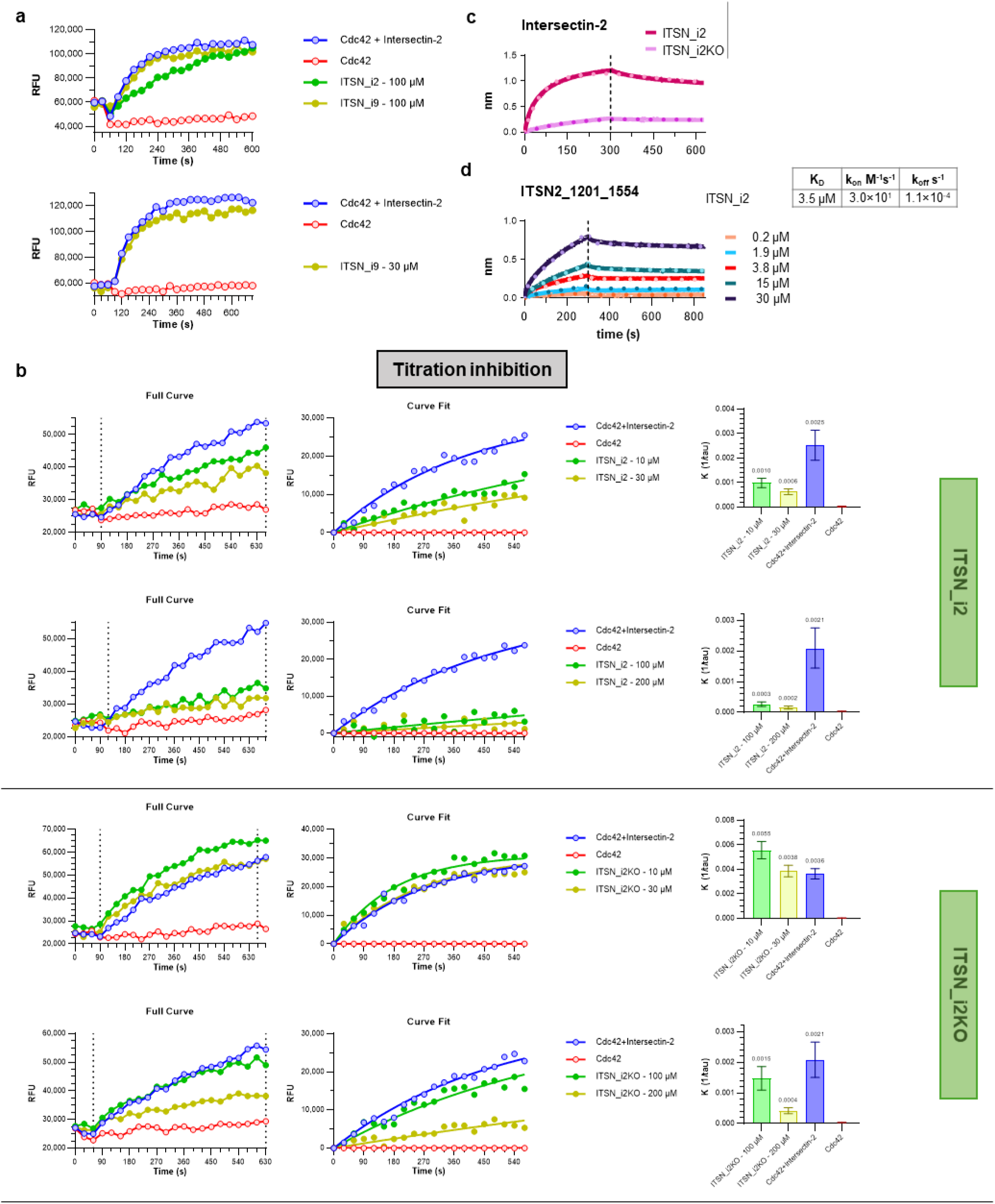
Binding and inhibition of Intersectin-2 by engineered inhibitors. a. Intersectin-2 inhibition by inhibitors Itsn1_i2 and Itsn1_i9 tested at 100 µM and 30 µM. b. Intersectin-2 inhibition by inhibitor Itsn1_i2 at different concentrations. c. BLI traces showing binding of Itsn1_i2 and its knockout variant. d. BLI traces to determine the kinetic parameters of Itsn_i2 interacting with recombinant Intersecting-2 residues 1201_1554. Binding parameters are summarized in the upper table.

**Supplementary Figure 11.**
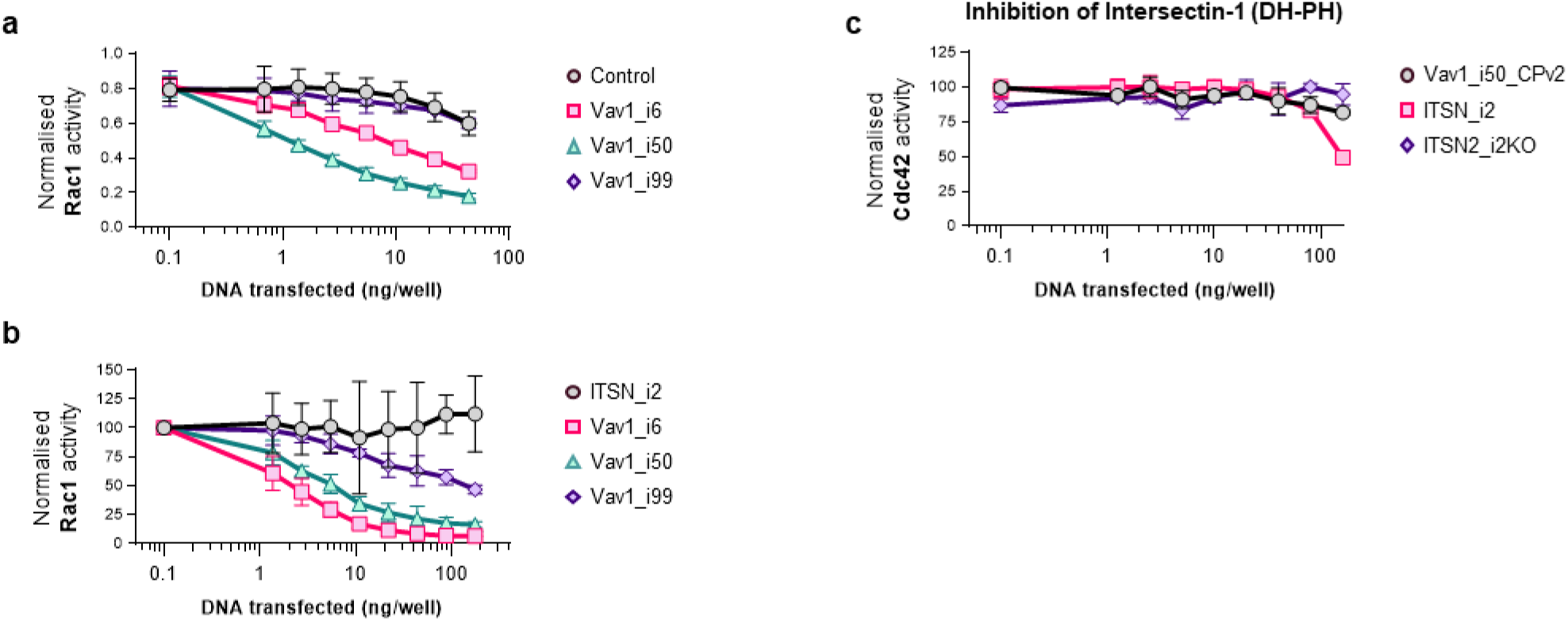
Selectivity of Intersectin inhibitor Itsn_i2 in cellular assays. a. Cellular assay showing Rac1 activity in cells co-transfected with Vav1 and increasing amounts of DNA encoding Vav1 inhibitors. Conditions include Vav1_ihn_6 (*pink*), Vav1_ihn_50 (*green*), Vav1_ihn_99 (*purple*), and no inhibitor control (*black*). b. Cellular assay showing Rac1 activity in cells co-transfected with Vav1 and increasing amounts of DNA encoding Vav1 inhibitors. Conditions include Vav1_ihn_6 (*pink*), Vav1_ihn_50 (*green*), Vav1_ihn_99 (*purple*), and ITSN2_i2 (*black*). c. Inhibition of Intersectin-1 mediated Cdc42 activation by Itsn_i2. Conditions include Intersectin-2 with ITSN2_i2 (*pink*), Itsn2_i2KO (*purple*), and Vav1_i50_CPv2 as a negative control.

**SUPPLEMENTARY TABLE 1.** List of inhibitors developed against Vav1 using motif grafting protocol in Rosetta.

| Design name | Parent scaffold (PDBid) | Scaffold origin | Size (#aa) | Mutations introduced | Epitope sequence | Yield (mg/L) | Protein MPNN | Modified name | Epitope sequence | Yield (mg/L) | Inhibition of Vav1 activity |
| --- | --- | --- | --- | --- | --- | --- | --- | --- | --- | --- | --- |
| 1ea3 | Influenza A virus | Influenza virus M1 protein | 153 | 22 | NSGADAIYLHLMAE | 2 | Yes | 1ea3_pMPNN_seq9 | AYADAIYLHLMLE | 9.3 | No |
| 1kdb | Staphylococcus aureus | Staphylococcal nuclease | 136 | 18 | LQAAEIYAWLMRL | 2 | No | n/a | n/a | n/a | n/a |
| 1nfn | Homo sapiens | Apolipoprotein E3 (APOE3) | 132 | 18 | TLAALIYDWLMLD | 0 | No | n/a | n/a | n/a | n/a |
| 1ng5 | Staphylococcus aureus | Staphylococcus aureus Sortase B | 211 | 20 | QAQFIYEKLMD | 10 (precipitated) | Yes | 1ng5_pMPNN_seq13 | KLQAQFIYEKLMD | 7.9 | No |
| 5efx | Homo sapiens | Rho guanine nucleotide exchange factor 2 | 144 | 13 | WIAAIIYSALMCP | 1.5 | Yes | 5efx_pMPNN_seq2 | KWIAAIIYAAIMRP | 8.4 | No |
| 5izw | Unidentified | PLS3-PPR | 101 | 16 | LAAAIYMIIMGIK | 0 | No | n/a | n/a | n/a | n/a |
| 6exu | Schizosaccharomyces pombe 972h- | Switch-activating protein 1 | 104 | 29 | EIVAEYIYELIMRLP | 10 (precipitated) | No | n/a | n/a | n/a | n/a |

**SUPPLEMENTARY TABLE 2.**
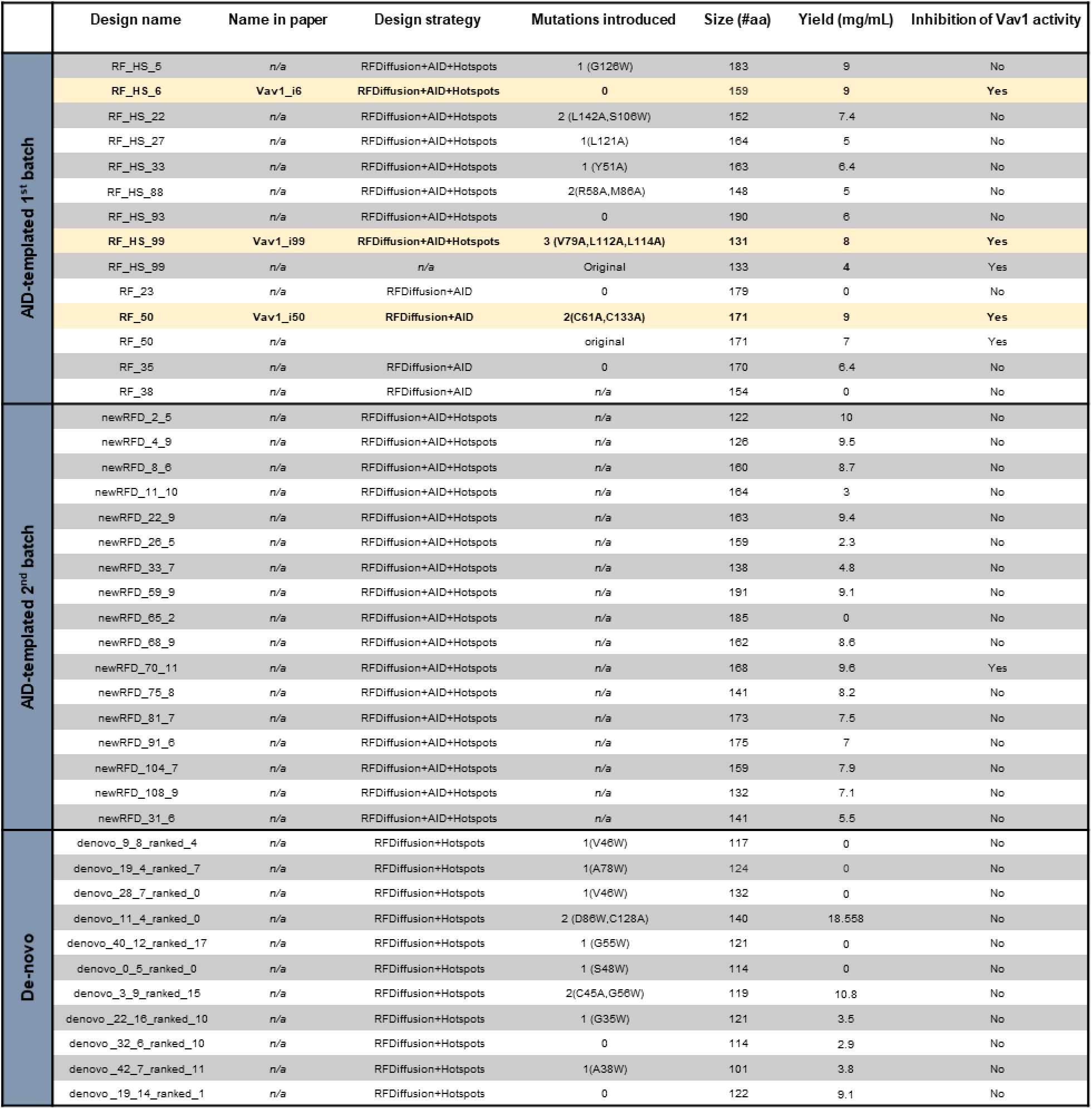
List of inhibitors developed against Vav1 using RFDiffusion. The first two groups are AID templated, while the third group is *de novo*.

**SUPPLEMENTARY TABLE 3.**
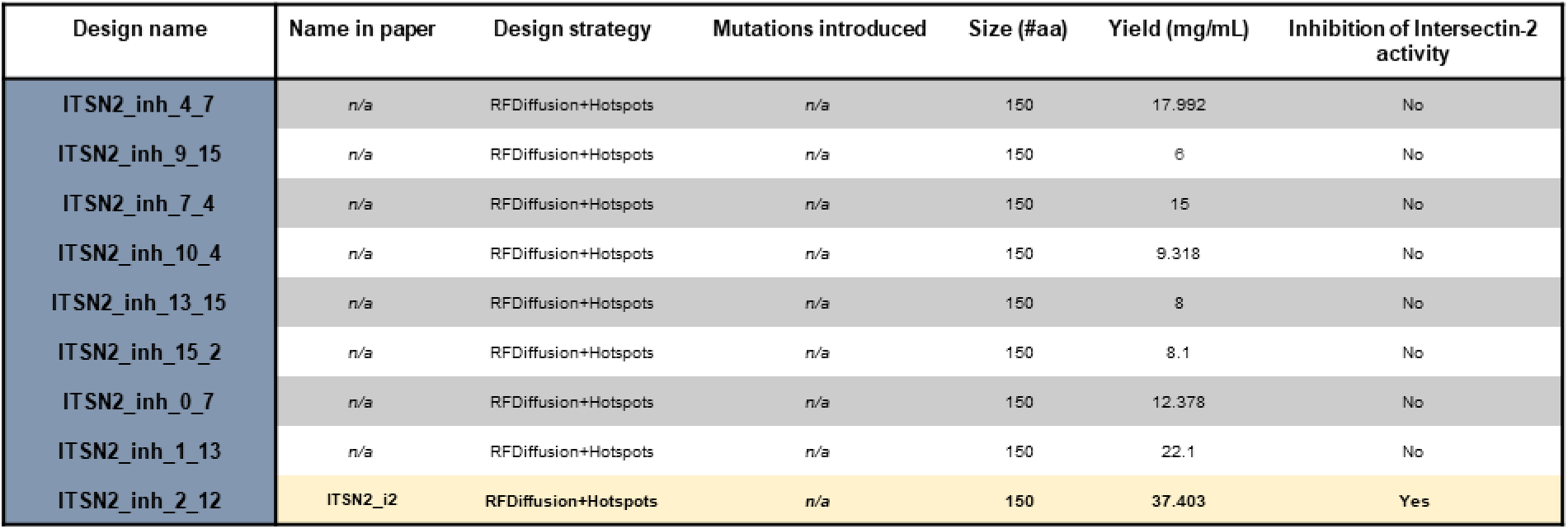
Inhibitors developed against Intersectin-2.

| Design name | Name in paper | Design strategy | Mutations introduced | Size (#aa) | Yield (mg/mL) | Inhibition of Intersectin-2 activity |
| --- | --- | --- | --- | --- | --- | --- |
| ITSN2_inh_4_7 | n/a | RFDiffusion+Hotspots | n/a | 150 | 17.992 | No |
| ITSN2_inh_9_15 | n/a | RFDiffusion+Hotspots | n/a | 150 | 6 | No |
| ITSN2_inh_7_4 | n/a | RFDiffusion+Hotspots | n/a | 150 | 15 | No |
| ITSN2_inh_10_4 | n/a | RFDiffusion+Hotspots | n/a | 150 | 9.318 | No |
| ITSN2_inh_13_15 | n/a | RFDiffusion+Hotspots | n/a | 150 | 8 | No |
| ITSN2_inh_15_2 | n/a | RFDiffusion+Hotspots | n/a | 150 | 8.1 | No |
| ITSN2_inh_0_7 | n/a | RFDiffusion+Hotspots | n/a | 150 | 12.378 | No |
| ITSN2_inh_1_13 | n/a | RFDiffusion+Hotspots | n/a | 150 | 22.1 | No |
| ITSN2_inh_2_12 | ITSN2_i2 | RFDiffusion+Hotspots | n/a | 150 | 37.403 | Yes |

**SUPPLEMENTARY TABLE 4.** Percent Vav1 inhibition of engineered inhibitors across a panel of different GEFs.

| GEF | Additional names | GTPase | % identity to Vav1<br>DH domain | Fold reduction ($K_{obs}$) | | |
| --- | --- | --- | --- | --- | --- | --- |
|  |  |  |  | Vav_i6 | Vav_i99 | Vav_i50 |
| Vav1 | n/a | Rac1 | 100 % | 73.4 | 16.2 | 16.9 |
| Vav3 | n/a | Rac1 | 61 % | 28.0 | 10.7 | 5.8 |
| Vav2 | n/a | Rac1 | 50 % | 1.8 | 1.5 | 1.3 |
| Tiam1 | n/a | Rac1 | 29 % | 1.7 | 1.3 | 0.9 |
| Trio | Triple functional domain protein | Rac1 | 23 % | 1.2 | 1.1 | 1.2 |
| Dock-10 | Dedicator of cytokinesis protein 10 | Rac1 | 14 % | 1.0 | 1.1 | 0.8 |
| LARG | ARHGEF12<br>Rho guanine nucleotide exchange factor 12 | RhoA | 26 % | 1.3 | 1.6 | 1.0 |
| p115 | ARHGEF1<br>RhoGEF | RhoA | 25 % | 2.1 | 2.1 | 1.1 |
| GEFH11 | ARHGEF11<br>Rho guanine nucleotide exchange factor 11 | RhoA | 25 % | 1.2 | 1.1 | 1.0 |
| GEFH1 | ARHGEF2<br>Rho guanine nucleotide exchange factor 1 | RhoA | 23 % | 0.9 | 1.1 | 1.3 |
| Intersectin-2 | n/a | Cdc42 | 25 % | 1.3 | 1.1 | 1.2 |

